# Cryo-mtscATAC-seq for single-cell mitochondrial DNA genotyping and clonal tracing in archived human tissues

**DOI:** 10.1101/2025.09.17.675534

**Authors:** Maren Salla, Benedikt Obermayer, Marie Cotta, Ekaterina Friebel, Juliana Campo-Garcia, Georgia Charalambous, Roemel Jeusep Bueno, Dustin Lieu, Patryk Dabek, Ashley Helmuth, George Tellides, Roland Assi, Katrin Bankov, Marco Lodrini, Hedwig Deubzer, Dieter Beule, Hattie Chung, Helena Radbruch, David Capper, Frank Heppner, Sarah C. Starossom, Caleb A. Lareau, Ilon Liu, Leif S. Ludwig

## Abstract

High-throughput clonal tracing of primary human samples relies on naturally occurring barcodes, such as somatic mitochondrial DNA (mtDNA) mutations detected via single-cell ATAC-seq (mtscATAC-seq). Fresh-frozen clinical specimens preserve tissue architecture but compromise cell integrity, thereby precluding their use in multi-omic approaches such as mitochondrial genotyping at single-cell resolution. Here, we introduce Cryo-mtscATAC-seq, a broadly applicable method for diverse pathophysiological contexts to isolate nuclei with their associated mitochondria (“CryoCells”) from frozen samples for high-throughput clonal analysis. We applied Cryo-mtscATAC-seq to the neurodegenerated human brain, glioblastoma (GBM), pediatric neuroblastoma, and human aorta, and implemented mitobender, a computational tool to reduce ambient mtDNA in single-cell assays. Our approach revealed regional clonal gliogenesis and microglial expansions in amyotrophic lateral sclerosis (ALS), persistence of oligodendrocyte progenitor cell (OPC)-like clones in GBM recurrence, mtDNA depth heterogeneity after neuroblastoma chemotherapy, and oligoclonal proliferation of smooth muscle cells in human aorta. In conclusion, Cryo-mtscATAC-seq broadly extends mtDNA genotyping to archival frozen specimens across tissue types, opening new avenues for investigation of cell state-informed clonality in human health and disease.

## Introduction

Flash-frozen tissue is one of the most widely archived biospecimens in clinical research, offering molecular preservation without chemical fixation. Thus, they are valuable for the longitudinal study of health and disease such as in the context of clinical trials, rare diseases, and tissues that are difficult to access prospectively, facilitating the assemblage of comprehensive and well-annotated clinical cohorts through international multicenter collaborations^1,2^. However, the disruption of the plasma membrane during freezing limits the recovery of intact cells, restricting the application of single-cell genomic applications largely to the profiling of the nuclear RNA and DNA content^3^.

In addition to the nucleus, mitochondria - an organelle central to cellular metabolism - harbor their own genome. The mitochondrial genome is compact (∼16.6 kb), has a high mutation rate, lacks protective histones, and replicates independently of the nuclear genome. It exists in multiple copies per cell, ranging from hundreds to thousands depending on the cell type. This multicopy nature gives rise to heteroplasmy, where wild-type and mutant mtDNA molecules can coexist within the same cell. Somatic mtDNA mutations accumulate during aging and are passed on to progenies in a non-Mendelian fashion^4–9^.

Single-cell genomic technologies have transformed our ability to resolve cellular heterogeneity at unprecedented resolution^10–12^. More recently, refined cell preparation methods enabled the parallel analysis of epigenetic states with mtDNA genotyping to leverage somatic mtDNA variants as naturally occurring barcodes to retrospectively infer clonal relationships in human cells. Compared to whole-genome sequencing-based clonal tracing, mtDNA-based approaches offer a more cost-efficient and scalable alternative that captures both cell type–specific chromatin states and clonal relationships from the same assay across tens to hundred thousands of cells^8,13–15^. Various studies demonstrated the utility of mtDNA variants in resolving clonal dynamics in humans. In the hematopoietic system, they have enabled clonally tracking the differentiation of stem cells into diverse blood lineages^13,14^ and tracing the clonal expansion and persistence of innate immune cells^16^. Applications also extend to solid tumors to reconstruct cancer evolution from dissociated tumor cells^17^ as well as spatially^18^ or tracking longitudinal clonal dynamics in blood malignancies^19,20^ highlighting the broad utility of mtDNA-based assays for investigating fundamental aspects of clonal architecture.

Modified cell isolation and library protocols enable capture of specific genomic features. Standard snATAC-seq recovers accessible nuclear regions via detergent washes^3^, whereas mtscATAC-seq, DOGMA-seq, and ASAP-seq use mild fixation and omit detergents to retain mitochondria during transposition^14,15,21^. To adapt these principles to fresh frozen tissue, we developed the isolation of Flash Frozen Cells’ (CryoCells) via thin cryosectioning and formaldehyde fixation to preserve mitochondrial–nuclear proximity for high-quality chromatin profiling with mtDNA genotyping. We validated this approach in murine spleen and applied it to a wide range of clinically relevant archived tissues - including *post mortem* human brain, primary and recurrent glioblastoma (GBM), pediatric tumors, and human vascular tissue - demonstrating its versatility across organ systems. To further account for ambient mtDNA, we developed mitoBender based on the popular method CellBender, which removes ambient RNA background in scRNA-seq data analysis^22^.

## Results

### The concept and isolation of ‘CyroCells’

We sought to develop a strategy to isolate single nuclei with their associated mitochondria for mtDNA genotyping from archived flash-frozen clinical specimens. Notably, current state-of-the-art protocols surrounding single-cell/nuclei sequencing from frozen samples rely on nuclei preparations, thereby largely losing cytoplasmic content, including mitochondria. Given that cells are intricately structured with cytoskeletal elements and internal membrane systems^23,24^, we reasoned that formaldehyde fixation to retain mitochondria within their respective cells as in mtscATAC-seq^13,14,21^ and DOGMA-seq^15^ could likewise be exploited to preserve cellular-like architecture in flash-frozen specimens^25^ (**Fig. 1a**). These approaches provided the foundation for adapting fixation and lysis conditions to frozen material, where detergent omission proved critical for mtDNA recovery^13–15,21^. By contrast, conventional ATAC-seq protocols rely on detergent-based washes tailored to nuclear DNA accessibility profiling^3^, which are incompatible with mitochondrial preservation. We therefore iteratively tested a variety of conditions to develop a workflow combining thin cryosectioning, controlled fixation, and mild lysis to yield “CryoCells” for downstream analysis (**Supplementary** Fig. 1). Key optimizations include: (i) cryosectioning tissue at 70 µm to reduce fixation time, improve uniformity, and allow region-specific sampling; (ii) careful control of cryostat temperature and direct collection of sectioned tissue into pre-cooled tubes, avoiding slide-melting procedures used in immunostaining; (iii) in-tube fixation along the tube wall to prevent scrolling and preserve morphology; (iv) use of a plastic pestle instead of glass douncers to minimize mechanical damage during disruption; (v) sequential filtration through 70 µm and 30 µm strainers with gentle syringe-assisted pressure to accommodate heterogeneous CryoCell sizes; and (vi) a low-speed centrifugation step to remove small debris and ambient mitochondria, overcoming the limitations of FACS (loss of mtDNA) and density gradients (fixation-induced density heterogeneity) (**Supplementary** Fig. 1). The resulting CryoCells were processed with the 10x Genomics scATAC-seq workflow, enabling mtDNA genotyping via mgatk and integration with chromatin accessibility profiles, consistent with mtscATAC-seq principles^13,14,21^.

**Figure 1.**
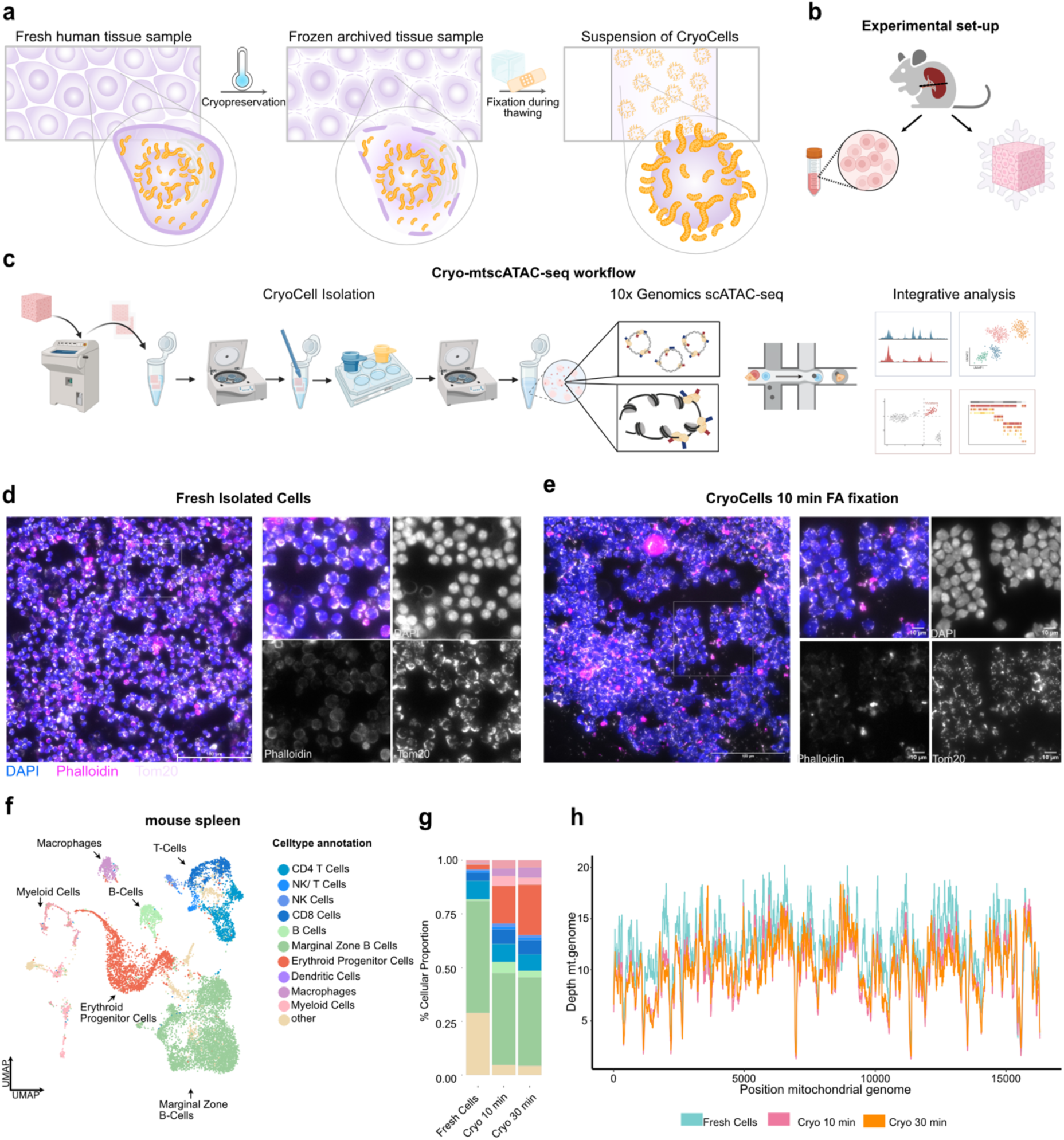
Benchmarking of fresh cryopreserved vs. flash-frozen mtDNA sequencing in murine spleen. **(a)** Concept of CryoCells. In fresh tissue, intact plasma membranes preserve nuclear–mitochondrial integrity. Flash freezing or cryopreservation disrupts membranes, whereas fixation during thawing stabilizes nuclei with their associated mitochondria, enabling CryoCell isolation. **(b)** Schematic of benchmarking workflow comparing CryoCells to freshly isolated spleen cells for mtDNA genotyping. **(c)** Isolation protocol for CryoCells from flash-frozen tissue. **(d,e)** Immunofluorescence staining of fresh spleen cells **(d)** and CryoCells **(e)** with DAPI (nuclei), Phalloidin (F-actin), and Tom20 (mitochondria). Scale bar, 100 µm. **(f)** UMAP embedding of fresh and CryoCell-derived spleen cells integrated by chromatin accessibility, demonstrating preservation of major immune cell lineages across preparation methods. **(g)** Cell type composition across conditions showing robust recovery of lymphoid, myeloid, and progenitor subsets. **(h)** Coverage profiles across the mitochondrial genome under fresh and fixed conditions.

### Benchmarking of fresh cryopreserved vs. ’Cryo’ mtDNA sequencing in murine spleen

To benchmark CryoCell isolation strategies, we bisected a spleen of a C57BL/6J mouse, with one-half being flash-frozen and stored at –70°C. Single-cell suspensions were generated right away from the other half following standard cell dissociation/isolation protocols and cryopreservation (**Fig. 1b,c** and **Methods**). For cryosectioning, the fresh-frozen tissue block was embedded in OCT, and equilibrated to –20°C before sectioning. Tissue scrolls were collected in precooled tubes for immediate processing or stored at –70°C for long-term preservation (**Fig. 1c**). For direct comparison, cryopreserved splenocytes were thawed and processed for mtscATAC-seq, while spleen CryoCells were isolated using the dedicated protocol outlined above and subjected to different fixation times (i.e., 10 min and 30 min; **Methods**). Importantly, mitochondrial retention was confirmed via immunostaining for the mitochondrial membrane protein Tom20 in both freshly isolated cells and formaldehyde-fixed CryoCells (**Fig. 1d,e**). All major splenic cell types were recovered across all three conditions following mtscATAC-seq, with differences in cellular composition likely reflecting differences of individual tissue sections and/or intrinsic biases of cell isolation protocols and/or sensitivity to freeze/thawing (**Fig. 1f,g** and **Supplementary** Fig. 2a,b). For example, CryoCells exhibited a higher fraction of erythroid progenitor cells, which may reflect the lack of red blood cell lysis when using cryosections and/or the inclusion of higher proportions of red relative to white pulp (**Fig. 1g**). Importantly, shallow sequencing revealed comparable mtDNA read depth and similar coverage patterns across all three conditions (**Fig. 1h** and **Extended Data** Fig. 1a-c). TSS enrichment was reduced in CryoCells relative to freshly isolated cryopreserved cells, but remained comparable to published single-nuclei ATAC-seq datasets^16,26,27^ (**Extended Data** Fig. 1d). mgatk analysis identified homoplasmic germline and selected heteroplasmic mtDNA mutations with mutational signatures being consistent with previous reports for mice and demonstrating the expected predominance of C>T transitions^28^ (**Extended Data** Fig. 1e,f). However, the number of heteroplasmic somatic mutations was relatively limited, consistent with the young age of the mice and previous reports showing a lower abundance of mtDNA mutations in mice relative to human samples^19,28^. Together, these data support the feasibility of mtDNA genotyping via mtscATAC-seq from flash-frozen specimens.

### mitoBender removes highly prevalent ambient mtDNA variants

Analogous to ambient RNA contamination in snRNA-seq, highly abundant mtDNA variants may contribute to ambient background signal in Cryo-mtscATAC-seq^22,29^ (**Fig. 2a**). While our prior work suggested ambient mtDNA levels to be low in 1% FA fixed cells in the original mtscATAC-seq implementation^14^, we reevaluated this in light of the use of flash frozen tissue. In principle, two scenarios can be distinguished. First, empty droplets may acquire ambient genomic material, yielding pseudobulk mtDNA variant proportions similar to those observed in bulk sequencing. Second, in droplets with CryoCells, the retained signal is dominated by its characteristic mtDNA variants of markedly higher heteroplasmy compared with empty droplets or droplets containing unrelated CryoCells (**Fig. 2a**). To address potential high-abundant mtDNA mixing, we extended principles from CellBender^22^ to develop mitoBender, which models background distributions of mtDNA allele counts from empty droplets and effectively reduces ambient signal (**Methods**).

**Figure 2.**
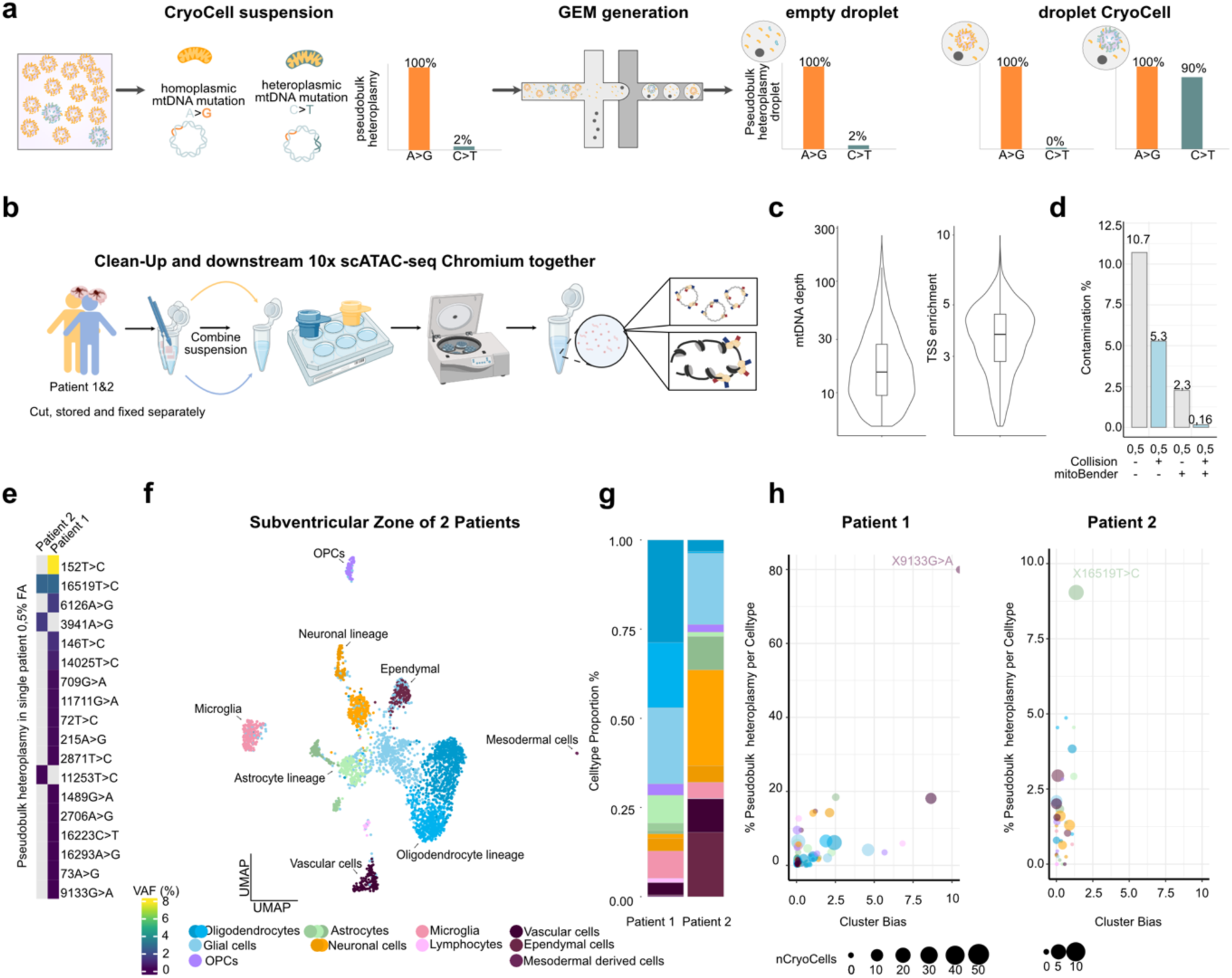
Removal of ambient mtDNA via mitoBender. **(a)** Schematic of ambient mtDNA distribution across the full dataset (far left), empty droplets (middle), and CryoCell-containing droplets (right), shown as pseudobulk heteroplasmy values of mtDNA variants. **(b)** Schematic overview of the experimental mixing setup. **(c)** Violin Plots of mtDNA depth and TSS enrichment as QC measurement of the data **(d)** Bar plot Contamination in %. Left (grey) and right (blue) barplots are derived from the same dataset. Grey from data without mitobender filter, whereas right (blue) mitobender filterd. Collision is based on mtDNA variants, which if taken out reduces the mixing. **(e)** Heatmap of high-confidence heteroplasmic variants detected in patient-specific mgatk calls from the 0.5% FA dataset. **(f)** UMAP embedding of CryoCells colored by cell type annotation. **(g)** Bar plot of cell type proportions per patient. **(h)** Bubble plot of cell type bias per patient based on potential clonally informative, high-confidence mtDNA variants. Point size reflects the number of CryoCells exceeding the variant threshold; y-axis denotes log₁₀ cluster bias scores.

We benchmarked mtDNA genotyping in human tissue and evaluated the feasibility of mtDNA background correction in a donor-mixing experiment using subventricular zone (SVZ) tissue from an ALS and a multiple sclerosis (MS) patient with distinct mtDNA haplotypes. Nuclear demultiplexing enabled the resolution of doublets (homotypic or heterotypic) and facilitated patient assignment of CryoCells^30–32^. In parallel, patient-specific homoplasmic germline mtDNA variants displayed a bimodal heteroplasmy distribution across cells - either nearly fixed (>99%) or absent (<1%) - in agreement with nuclear genotyping (**Extended Data** Fig. 2a,b). Unlike nuclear doublets, mtDNA background arose from collisions, in which mtDNA molecules (ambient mtDNA) from the other donor were co-encapsulated within droplets.

After stringent QC filtering (mtDNA coverage > 5 and TSS enrichment > 1.5; **Fig. 2c**) and mitoBender correction, ambient mtDNA contamination was effectively reduced to 0.14%, comparable to the original mtscATAC-seq implementation^14^ (**Fig. 2d** and **Extended Data** Fig. 2a,b).Importantly, clonally informative somatic mtDNA variants are typically those with high variant allele frequency (VAF) in individual CryoCells but low overall pseudobulk abundance (<1–2% heteroplasmy in highly polyclonal systems)^14,33^. Restricting variant calling to CryoCells confidently assigned to one donor revealed unique variants per patient at this low pseudobulk heteroplasmy, with the exception of a single shared variant (**Fig. 2e**). For example, mt.16591T>C was detected in both donors but lies within the mutationally prone D-loop region^34^ (**Fig. 2e**). We further detected all major CNS-resident cell types, with ependymal cells uniquely present in the SVZ, indicating that nuclear chromatin accessibility profiles from fixed samples can reliably capture cellular identity (**Fig. 2f,g**). Cell-bias analysis of clonally informative variants showed variable pseudobulk heteroplasmy across cell types with in particular one variant being restricted to mesoderm-derived cells, suggestive of an early embryonic origin (**Fig. 2h**). Together, these results demonstrate that mitoBender effectively removes ambient signal from highly abundant mtDNA variants, such as those from germline origin when multiplexing multiple donors. In contrast, clonally informative variants remain largely unaffected by such mixing, as their low abundance in pseudobulk prevents misclassification, enabling their robust use as somatic clonal markers in complex tissues. This is analogous to the relatively low contribution of lowly expressed genes to ambient RNA in scRNA-seq analysis. Building on these validations, we applied Cryo-mtscATAC-seq across multiple tissues to demonstrate its broader applicability.

### mtDNA genotyping reveals gliogenesis and clonal microglia dynamics in human nervous system pathologies

Genetic barcoding in mice and the whole genome sequencing (WGS)-based detection of somatic mutations have provided foundational insights into the polyclonal origin and phylogenies of ectodermal-derived CNS resident cells^35–37^. To evaluate the potential of Cryo-mtscATAC-seq for investigating aspects of clonality in the human brain, we analyzed archived fresh-frozen clinical specimens from an 80-year-old male patient with amyotrophic lateral sclerosis (ALS). ALS is characterized by a progressive disease course affecting upper and lower motor neurons of different brain regions^38^. The selection of brain regions were guided following neuropathologic assessments to reflect regional and disease heterogeneity, focusing on four anatomically and functionally distinct areas: the prefrontal cortex (PFC) serving as a patient-internal control that is typically not prominently affected in ALS, the motor cortex (1MNC), the medulla oblongata (MO), and the cervical spinal cord (CSC) (**Fig. 3a,b**). Following stringent filtering and doublet removal using Amulet^30^, 13,699 CryoCells were recovered across all four brain regions. As expected, cortical and non-cortical regions displayed distinct cellular compositions, with excitatory and GABA inhibitory neurons being enriched in cortical areas (PFC and 1MNC)^39–41^ (**Fig. 3b,d** and **Supplementary** Figure 3a-c). Furthermore, region-specific glial cell compositions/frequencies were observed, highlighting the differences between regions, including their expected depletion in the medulla oblongata, and high scATAC-seq data quality enabling the differentiation of oligodendroglial (OL) subtypes. Notably, microglial cellular proportions correlated with disease manifestation, showing their increasing abundance from cortical regions to the cervical spinal cord (**Fig. 3b,d** and **Extended Data** Fig. 3a-c). As under physiological conditions, microglia are typically evenly distributed throughout the CNS^42^, this suggests local inflammatory remodeling, including increased T-cell infiltration in the medulla oblongata and the cervical spinal cord, consistent with previous reports of inflammatory microglial states in ALS^43^.

**Figure 3.**
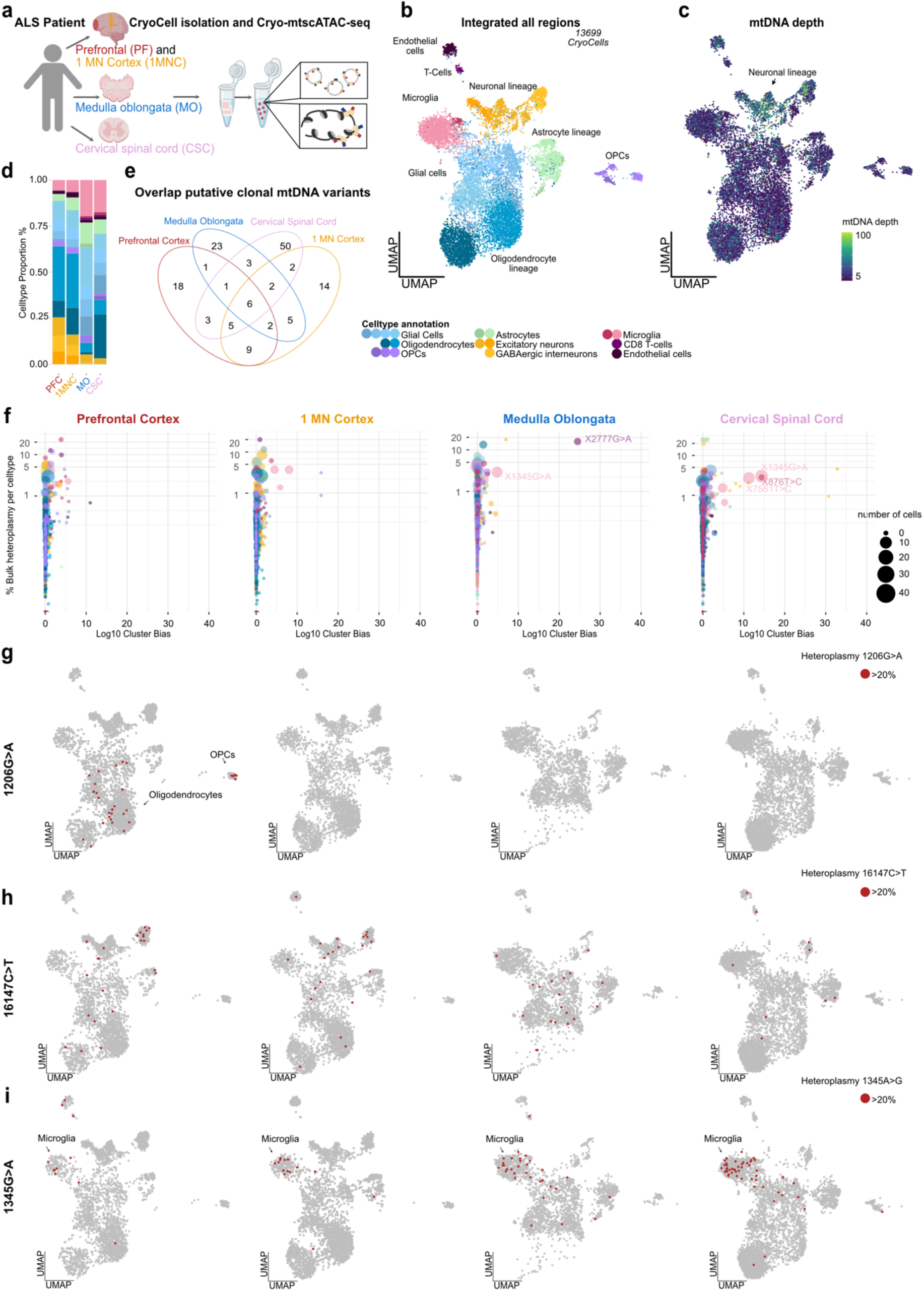
Region-specific mitochondrial clonal architecture in CNS tissue from an ALS patient. **(a)** Schematic overview of Cryo-mtscATAC-seq applied to four anatomically distinct CNS regions from a single ALS patient: prefrontal cortex, 1 motor cortex, medulla oblongata, and cervical spinal cord. **(b)** UMAP embedding of 13,699 CryoCells integrated across the four regions and annotated by cell type based on chromatin accessibility. **(c)** UMAP colored by per-cell mtDNA depth, highlighting cell type-specific variation. **(d)** Bar plot of cell type composition across the four regions. **(e)** Venn diagram of informative heteroplasmic mtDNA variants detected across regions (filters: n_cells_conf_detected ≥ 5, strand correlation > 0.65, log₁₀(VMR) > –2, mean coverage ≥ 5). **(f)** Bubble plots depicting lineage bias analysis per region, showing –log₁₀(adjusted Kruskal–Wallis p-value) cluster bias versus mean pseudobulk heteroplasmy per celltype for variants with >20% heteroplasmy. Point size reflects the number of CryoCells carrying a unique mtDNA variant (left to right: PFC, 1MNC, MO, CSC). **(g)** UMAP projection of cells with >20% heteroplasmy (red) for the region-restricted mt.1206G>A clone in oligodendrocyte-lineage cells of the prefrontal cortex. **(h)** UMAP projection of cells with >20% heteroplasmy (red) for mt.16147C>T, present across all cell types and regions. **(i)** UMAP projection of cells with >20% heteroplasmy (red) for the microglia-enriched mt.1345G>A clone spanning all regions.

Using **mgatk**, we identified 2,586 unique somatic mtDNA variants across all brain regions, which were distributed throughout the mitochondrial genome, with consistent coverage patterns being observed across all anatomical regions (**Extended Data** Fig. 4a,b**,e** and **Methods**). Analysis of mutational signatures showed the expected abundance of C>T and T>C transitions (**Extended Data** Fig. 4c), corroborating robust mutation calling from flash-frozen human tissue^14^. Neuronal CryoCells exhibited higher mtDNA depth compared to other cell types (**Fig. 3c**), likely reflecting their elevated energy demands^44^ and high mitochondrial abundance in the soma of neuronal cells^45^. mgatk analysis revealed 37 homoplasmic mtDNA variants shared across all regions, reflecting their germline origin (**Extended Data** Fig. 4a). Filtering for clonally informative variants identified 127 heteroplasmic mtDNA variants, including six variants shared across all regions (**Fig. 3e**). However, most somatic mtDNA variants were region- and cell type-specific suggesting they arose locally or presenting region-specific developmental processes (**Fig. 3e** and **Extended Data** Fig. 4d), as well as highlighting the polyclonal nature of the human brain^35–37,46^.

A more granular evaluation of individual mtDNA variants (**Fig. 3f** and **Extended Data** Fig. 5d) revealed cell type-specific biases, including within one anatomical region. For example, mt.1206G>A was exclusively present in CryoCells of the oligodendrocyte lineage in the prefrontal cortex (**Fig. 3g** and **Extended Data** Fig. 4g), along with additional oligodendrocyte-specific variants being prevalent in other regions (**Extended Data** Fig. 4h), in line with genetic fate-mapping studies demonstrating their regional oligoclonal expansion in mice^37^. The presence of mt.1206G>A in both OPCs and mature OLs (**Fig. 3g**), but regional restriction to the prefrontal cortex, further supports the notion of ongoing local and clonal oligodendrogenesis contributing to myelination plasticity throughout life^47,48^. In contrast, more broadly shared mtDNA variants such as mt.16147C>T may have originated during early embryonic development (**Fig. 3h**), given their presence across diverse cell type populations^6,49^.

Given the utility of mtDNA mutations to track clonality in innate immune cells^16^, we closely examined the brain’s resident immune cells, microglia, for evidence of clonal expansion. Subclustering of microglia revealed no region-specific phenotypic clusters, yet differential chromatin accessibility analysis uncovered regionally enriched peaks, including increased *PCDH9* accessibility in prefrontal cortex microglia and *NAV3* in the spinal cord, consistent with known region-specific and neurodegeneration-associated signatures^39,50^ (**Extended Data** Fig. 3a,b**,d**). Moreover, microglia harbored region-specific mtDNA variants, with at least one unique clone being identifiable per region (**Extended Data** Fig. 3e,f). Notably, we recovered the largest number of distinct microglia clones in the cervical spinal cord (**Fig. 3i** and **Extended Data** Fig. 3e,f). In the other regions, less clones defined by one unique variant could be identified suggesting oligoclonal expansions rather than dominant monoclonal responses (**Extended Data** Fig. 3e,f). Interestingly, we found one variant mt.1345G>A to be shared among microglia across regions with increasing prevalence from cortical regions to the cervical spinal cord, coinciding with a higher proportion of microglia in these areas (**Fig. 3i** and **Extended Data** Fig. 3e,g).

Chromatin accessibility analysis identified *HOXA5* as a differentially accessible locus in the mt.1345G>A clone (**Extended Data** Fig. 3h,i), a key regulatory transcription factor of myelopoiesis^51^. We also identified the region- and cell type-specific mt.2777G>A mutation to mark a unique T cell clone within the medulla oblongata (**Fig. 3f**). These results align with previous findings of microglia-driven inflammatory reactions in ALS^52^, and fate mapping studies in mice describing microglia as long-lived, self-renewing tissue-resident myeloid cells, which exhibit region-specific multiclonality^53^ likely originating from spatially restricted progenitor niches^42^. Along these lines, mtDNA-based tracing supports the regional persistence of microglia clones in adulthood, and validates Cryo-mtscATAC-seq as a robust tool for studying regional and cell type-specific clonal dynamics in the human CNS.

### Clonal dynamics in human glioblastoma evolution

Clonal dynamics are of fundamental interest to dissect tumor evolution, and prior work has demonstrated the utility of somatic mtDNA mutations to mirror longitudinal disease history and resolve heterogeneity in human blood malignancies^19,20,54^. To evaluate the potential of Cryo-mtscATAC-seq in capturing tumor dynamics of solid cancers within the CNS, we analyzed human glioblastoma (GBM) samples. GBM is characterized by rapid, diffuse growth, high heterogeneity, and the processes underlying its origins and clonal evolution during disease progression remain unclear. GBM has been shown to consist of highly heterogeneous and plastic cell states that resemble diverse neural cell lineages including neurodevelopmentally restricted cell types such as outer radial glia of the SVZ, a neural stem cell niche^55–58^.

Specifically, we applied Cryo-mtscATAC-seq to archived matched primary and recurrent tumors from two GBM patients (**Fig. 4a,b** and **Methods**). We annotated cell clusters using marker genes and published gene expression signatures/metaprograms derived from scRNA-seq data. Annotations were based on their predominant features, revealing astrocytic (AC)-like, NPC-like, mesenchymal-like, and oligodendrocyte progenitor cell (OPC)-like states (Fig. 4b,d, **Extended Data** Fig. 5a-c)^55,56,59^. Non-malignant populations such as microglia, tumor-associated macrophages (TAMs), endothelial cells, pericytes, and T cells were readily identified (**Fig. 4b,d** and **Extended Data** Fig. 5b,c). Mitochondrial genotyping revealed expected mutational signatures^14^ (**Extended Data** Fig. 6a-c), and an increased mtDNA depth was consistently observed in AC-like phenotypes, as well as endothelial cells across patient samples (**Fig. 4c** and **Extended Data** Fig. 6d-j). Cellular state abundances varied across patients and between primary and recurrent tumors (**Fig. 4d**), underscoring pronounced intra- and interpatient heterogeneity in GBM.

**Figure 4.**
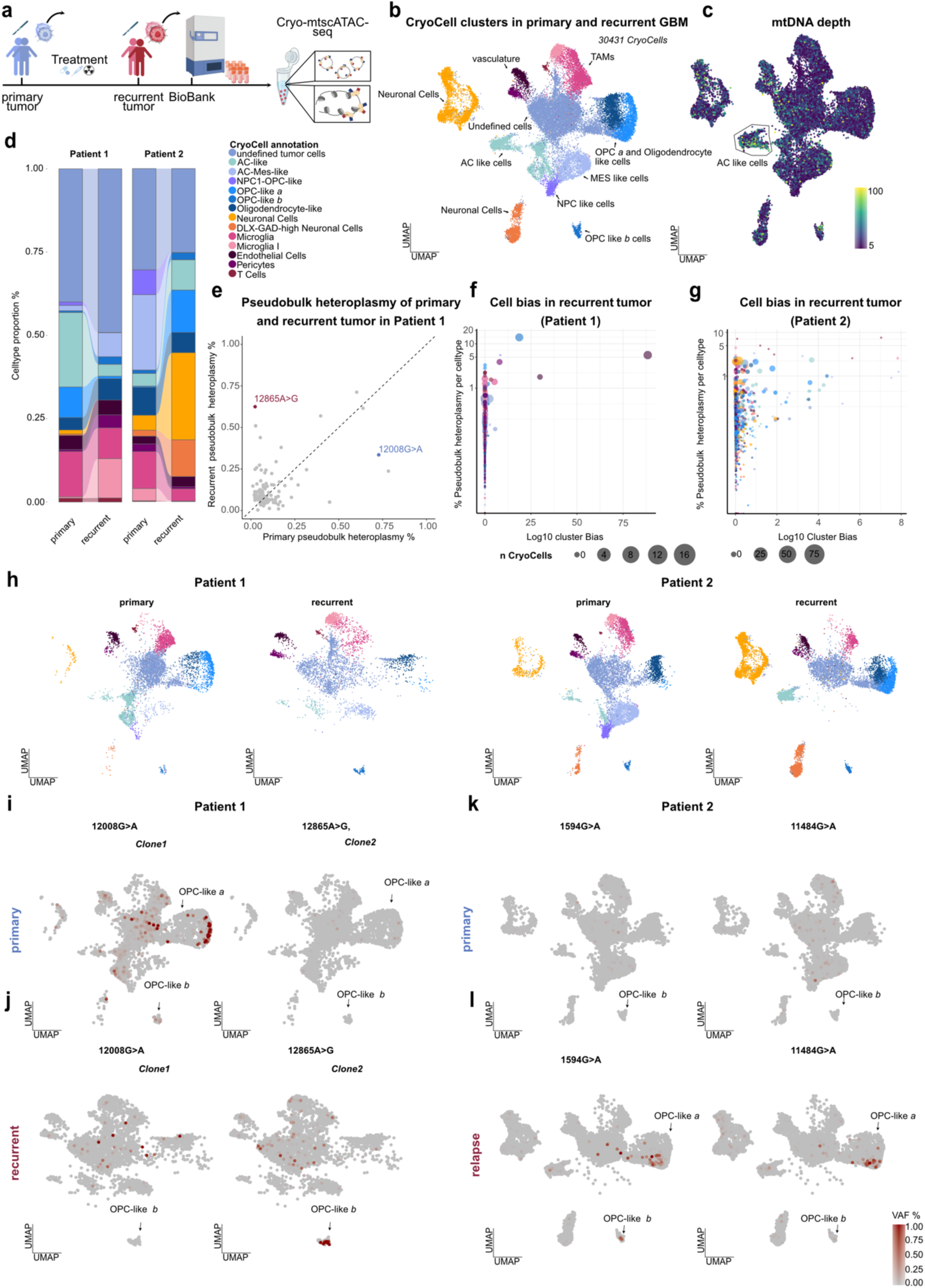
Clonal dynamics between primary and recurrence GBM. **(a)** Overview of tumor sampling and Cryo-mtscATAC-seq for paired primary and recurrence tumors from two GBM patients. **(b)** UMAP embedding of 30,431 CryoCells from all four datasets, colored by cell identity. **(c)** UMAP embedding colored by per-cell mtDNA depth. **(d)** Alluvial plot of cell type proportions in primary and recurrence tumors of each patient. **(e)** Scatterplot comparing bulk heteroplasmy levels of informative mtDNA variants between primary and recurrence tumors of patient 1. **(f)** Bubble plot showing pseudobulk heteroplasmy per cell type for mtDNA variants in the recurrence tumor of patient 1. Cells harboring >20% heteroplasmy are displayed with point size reflecting the number of cells; y-axis shows adjusted log₁₀ cluster bias scores, highlighting variant-specific cell type associations. **(g)** Bubble plot showing pseudobulk heteroplasmy per cell type for mtDNA variants in the recurrence tumor of patient 2, as in (f). **(h)** UMAP projections of CryoCells per dataset, colored by cell identity, split for each sample highlighting different celltype compositions per sample. **(i–l)** UMAP projections highlighting the distribution of cells harboring selected mtDNA variants (allele frequency scale: 0 = grey, 1 = red). (**i,j)** mt.12008G>A and mt.12865A>G in patient 1, shown for the primary (**i**) and recurrent (**j**) tumor. (**k,l)** mt.1594G>A and mt.11484G>A in patient 2, shown for the primary (**k**) and recurrent (**l**) tumor.

Longitudinal dynamics were resolved by focusing on clonally informative mtDNA variants shared between primary and recurrent tumors (**Methods**). In patient 1, pseudobulk analysis revealed a marked shift in mtDNA variant frequencies, with clone 1 (mt.12008G>A) enriched in the primary tumor (0.78% pseudobulk heteroplasmy) and clone 2 (mt.12685A>G) predominating at recurrence (0.62%) but already detectable at low levels in the primary tumor (0.024%) (**Fig. 4e**). Strikingly, this directed attention to two distinct OPC-like clusters (termed OPC-like-a and OPC-like-b) (**Fig. 4b** and **Extended Data** Fig. 5a-c**, f,g; Extended Data** Fig. 7a): OPC-like *a* cells clustered near oligodendrocyte-like populations in the UMAP, while OPC-like *b* cells formed a more distinct cluster and co-engaged NPC-associated programs (**Extended Data** Fig. 5a–c**,f,g**). Gene activity and GO analyses further distinguished OPC-like *a* cells from OPC-like *b* populations, showing that OPC-like-a cells retained differentiation-associated programs, while OPC-like-b cells were enriched for extracellular matrix remodeling processes (**Extended Data** Fig. 7b,c). At the regulatory level, only OPC-like *a* showed increased chromatin accessibility at canonical OPC transcription factors *OLIG2* and *SOX10* activity^60^ (**Extended Data** Fig. 7d,e), as well as stemness-associated factors SOX2 and JUN^58^ (**Extended Data** Fig. 7f,g). By contrast, OPC-like *b* cells lacked *SOX10* and *SOX2* activity but showed *MYT1* to be more accessible, a regulator of oligodendrocyte differentiation aberrantly activated in GBM^61^ (**Extended Data** Fig. 7h). Both OPC-like states additionally exhibited *BACH1* activity (**Extended Data** Fig. 7i), a transcription factor linked to temozolomide resistance and peritumoral OPC-like infiltration^56,62^.

Finally, mapping clonal variants onto these states revealed clone 1 enriched in OPC-like *a* cells of the primary tumor, while clone 2 marked OPC-like *b* cells at recurrence (**Fig. 4f,i,j**). Correspondingly, OPC-like *a* cells were strongly depleted at recurrence (primary: 9.16%, relapse: 0.076% of CryoCells), whereas OPC-like *b* cells expanded (primary: 0.052%, relapse: 2.28%) (**Fig. 4d,h**), suggesting that mt.12685A>G bearing CryoCells survived treatment, expanded, and transitioned into a more aggressive state, while mt.12008G>A-marked CryoCells were lost. In patient 2 however, mt.1594G>A and mt.11484G>A were enriched at recurrence across OPC-like *a* populations (pseudobulk heteroplasmy of all CryoCells: 0.1% each). Both variants were detectable at low heteroplasmy in the primary tumor (mt.1594G>A, 0.02%; mt.11484G>A, 0.05%), indicating a 2–5-fold clonal expansion of OPC-like *a* populations upon recurrence (**Fig. 4g,k,l**). Together, these results highlight the ability of Cryo-mtscATAC-seq to resolve clonal expansions and divergent functional programs underlying OPC-like states during GBM evolution and recurrence.

### mtDNA copy number heterogeneity in pediatric tumors

To evaluate the applicability of Cryo-mtscATAC-seq to pediatric tumors, we profiled neuroblastoma (NB), an aggressive embryonal tumor of the sympathoadrenal lineage, notable for its rather low mutational burden, clinical heterogeneity and frequent therapy resistance^63–65^. Despite its clinical relevance, the molecular characteristics remain incompletely understood, partly due to limited investigation at single-cell resolution and access to fresh tissue samples. We first analyzed two NB specimens from the same two-year-old female patient, collected longitudinally—one at initial diagnosis (NB01) and a second at a subsequent “second-look” surgery (NB02) separated by 8 months of treatment (**Fig. 5a**). Following quality control, 13,982 CryoCells passed filtering (**Extended Data** Fig. 8a,b), capturing the major cell populations typically observed in NB^66,67^, including a prominent neuroendocrine tumor compartment (**Fig. 5b**). Cell type annotations based on chromatin accessibility profiles aligned well with known lineage markers^66,68^ (**Extended Data** Fig. 8c, top). Notably, mtDNA coverage was elevated in tumor cells compared to non-tumor populations, suggesting an increased mitochondrial-energetic demand (**Fig. 5c**). We further observed variation in mtDNA depth across tumor clusters and a notable reduction in the “second-look sample” (**Fig. 5c** and **Extended Data** Fig. 8d), highlighting the utility of Cryo-mtscATAC-seq to investigate aspects of mitochondrial copy number variation and heterogeneity in complex tissues. While the average mtDNA coverage per tumor cell exceeded 30× (**Fig. 5d**), only five clonally informative somatic mtDNA variants in the primary tumor and six in the second-look specimen were detectable (**Extended Data** Fig. 8e), also suggesting a low somatic mtDNA mutational burden of NB. Further, pseudobulk heteroplasmy analysis revealed three mtDNA variants with bias towards one of the timepoints (e.g., 11825G>A), indicating dynamic clonal shifts over the treatment course (**Fig. 5e,f**). Lineage-resolved analysis uncovered cell type-specific variant enrichment in the endothelial cell compartment highlighted by mt.953T>C, suggesting lineage-specific somatic mtDNA evolution within the tumor microenvironment (**Fig. 5f**).

**Figure 5.**
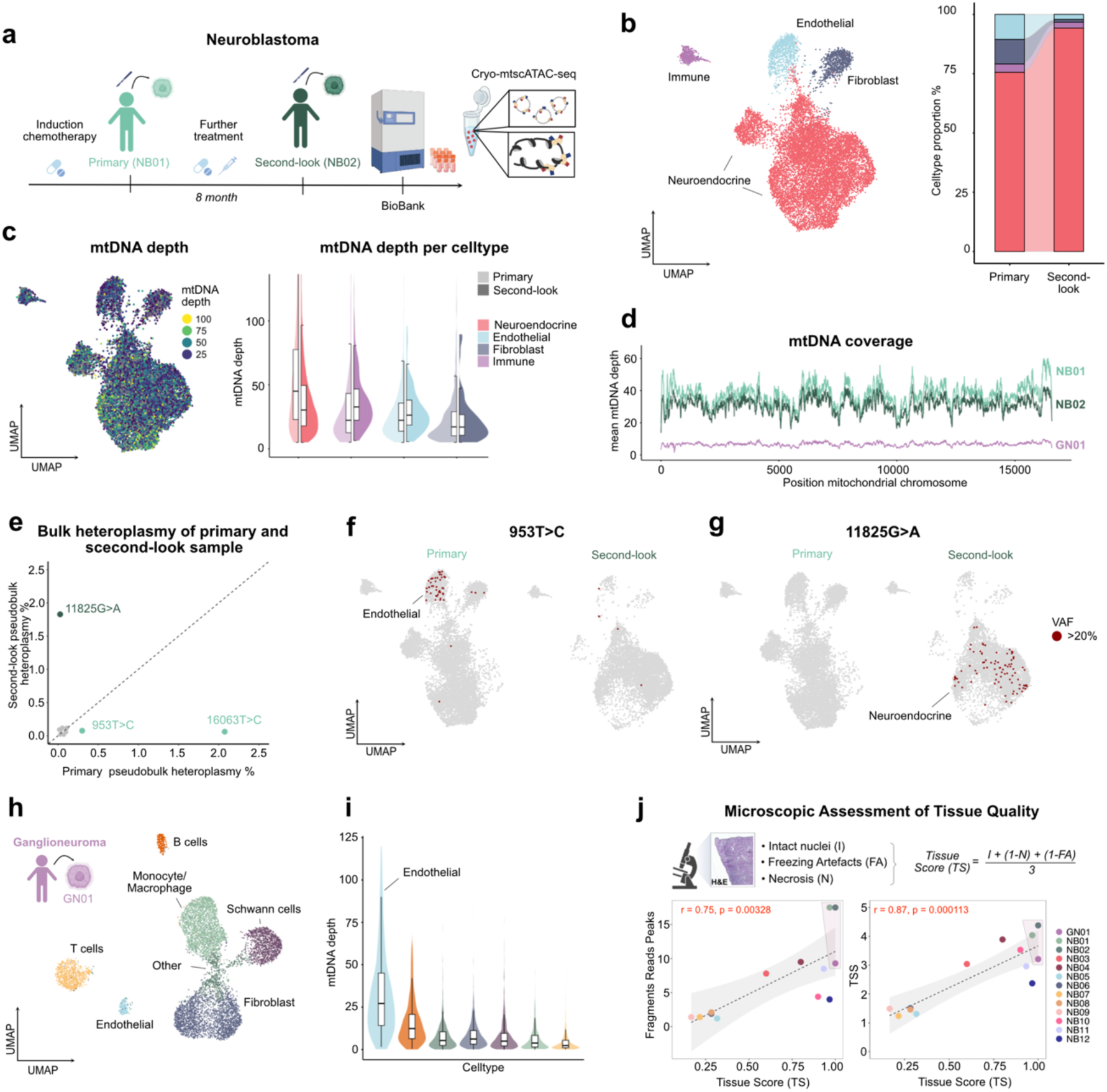
Somatic mtDNA variant detection in pediatric neuroblastic solid tumors using Cryo-mtscATAC-seq outside the CNS. **(a)** Schematic of sample collection and Cryo-mtscATAC-seq for paired primary (NB01) and second-look (NB02) surgical resections from a high-risk neuroblastoma patient. **(b)** UMAP embedding of 13,982 CryoCells from both time points, colored by cell identity (left); alluvial plot showing temporal shifts in cell type composition (right). **(c)** UMAP representation of per-cell mtDNA read depth, shown as a continuous color scale (left). Violin plots of mtDNA read depth per cell, stratified by cell type and sample identity (right). **(d)** Mitochondrial positional coverage profiles color coded by datasets **(e)** Scatterplot comparing pseudobulk heteroplasmy levels of informative mtDNA variants between primary and second-look surgery tumor samples of the high-risk neuroblastoma patient **(f,g)** UMAP projection of region-specific mtDNA variants in neuroendocrine and endothelial cells, showing the distribution of mt.953T>C (f) and mt.11825T>C (g) across time-points (primary tumor, left; second-look tumor, right). **(h)** UMAP embedding of 8,327 CryoCells from a ganglioneuroma patient profiled via Cryo-mtscATAC-seq, colored by cell identity **(i)** Violin plot showing per-cell mtDNA depth by cell type **(j)** Description of the tissue score (TS) calculation and scatter plots correlating TS with scATAC-seq quality control metrics (fragments in peaks and TSS enrichment) across 12 neuroblastoma and one ganglioneuroma sample.

We further profiled a ganglioneuroma (GN01), a benign, neural crest-derived peripheral neuroblastic tumor characterized by terminal neuronal differentiation^64^, from a 6.7-year-old patient. Cryo-mtscATAC-seq resolved its cellular composition, revealing fibroblasts, Schwann cells, endothelial and different immune cell populations^69^ (**Fig. 5g** and **Extended Data** Fig. 8c). While overall sequencing depth was lower, 14 clonally informative mtDNA variants recovered in more than five CryoCells (ncof>5) were identified, consistent with the additional age-related somatic accumulation (**Extended Data** Fig. 8e,f**,g**). Notably, mtDNA depth was highest in endothelial cells (**Fig. 5i**).

Over the course of our primary tumor profiling efforts, we further observed a strong link between histological tissue quality and genomic data quality. To systematically capture this, we developed a microscopic scoring system based on haematoxylin and eosin stain-stained (H&E) sections (**Fig. 5h**). Across 13 biobanked pediatric tumor samples (12 neuroblastoma, 1 ganglioneuroma), the score well-correlated with key scATAC-seq quality metrics, including the number of fragments in peaks (r = 0.75, p < 0.01) and TSS enrichment (r = 0.87, p < 0.01) (**Fig. 5h**), highlighting its utility as a practical predictor of data quality in Cryo-based single-cell chromatin accessibility profiling.

### Clonal tracing of smooth muscle cells in human aorta

Clonal expansion, reprogramming, and transdifferentiation of smooth muscle cells (SMCs) have been implicated across vascular pathologies. Yet, current evidence primarily stems from murine models and human X-inactivation studies^70–75^. Single-cell isolation from vascular tissues poses a major challenge due to structural heterogeneity and, in disease contexts, due to calcification and extensive extracellular matrix remodeling. In particular, SMCs are highly susceptible to enzymatic digestion, which compromises cell viability and recovery during tissue dissociation^76^. Given the central role of epigenomic regulation in SMC state transitions^77^, we assessed the utility of Cryo-mtscATAC-seq to characterize the proximal and mid-ascending part of the aorta from a 37-year-old male with a root- dominant thoracic aortic aneurysm (**Fig. 6a**).

**Figure 6.**
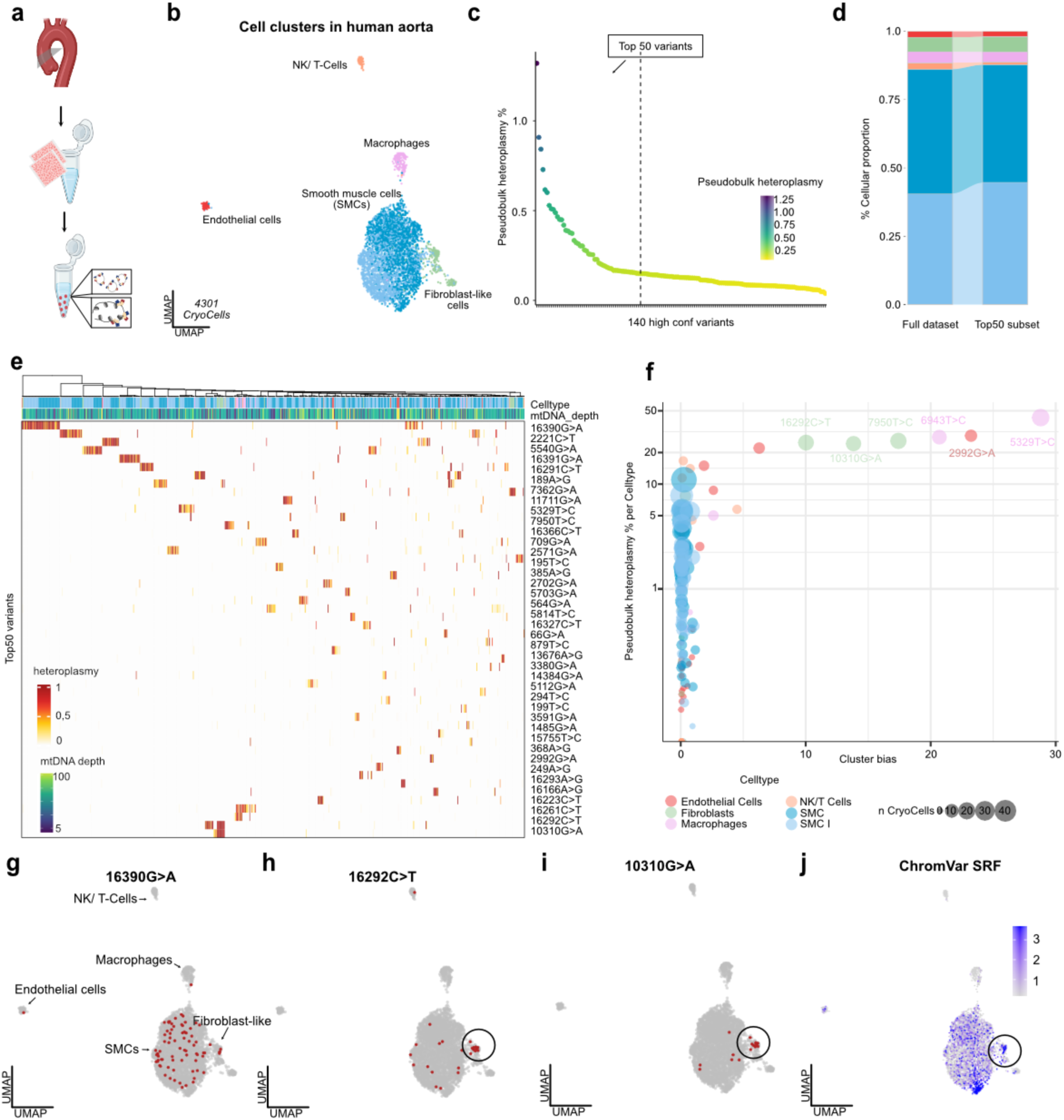
Mitochondrial variant profiling and lineage bias in human aortic tissue. **(a)** Schematic experimental overview. **(b)** UMAP embedding of 6,531 CryoCells derived from human aortic tissue, colored by annotated cell type. **(c)** Pseudobulk Heteroplasmy of 140 high confident mtDNA variants detected in the whole dataset. **(d)** Alluvial plot of cell type proportions in the full dataset and in the subset of CryoCells carrying at least one of the top 50 variants with >20% VAF. **(e)** Heatmap of heteroplasmy across the top 50 informative mtDNA variants. Each column represents a CryoCell, hierarchically clustered by variant abundance. **(g)** Bubble plot of pseudobulk heteroplasmy levels for mtDNA variants with >20% heteroplasmy in at least one cell type. Point size indicates the number of cells above threshold, and y-axis denotes log₁₀ cluster bias scores, highlighting variant-specific enrichment. **(h–j)** UMAP projections of distinct clones defined by mtDNA variants (>20% VAF; red = mutant cells, grey = others): (h) mt.16390G>A, (i) mt.16292C>T, and (j) mt.10310G>A. **(k)** ChromVAR-inferred SRF activity projected on UMAP.

Following quality control (e.g., >1,000 nuclear fragments, TSS enrichment >2), we retained 4,301 high-quality CryoCells spanning all major vascular cell types (**Extended Data** Fig. 9a), with SMCs constituting the most abundant population - consistent with previous scRNA-seq and snATAC-seq datasets^77^ (**Fig. 6b**). Endothelial cells displayed the highest mtDNA read depth among vascular populations (**Extended Data** Fig. 9b). Mitochondrial genotyping identified 140 high-confidence heteroplasmic variants, with pseudobulk heteroplasmy levels ranging from 0.039% to 1.3% suggesting most clones to be small in size (**Fig. 6c**). Variants were distributed throughout the mitochondrial genome, with an apparent enrichment in the D-loop region - consistent with its previously described mutation hotspot^34^ (**Extended Data** Fig. 9c,d). Mutational co-occurrence within single CryoCells was rare (**Fig. 6e**), and the mutation spectrum was consistent with replication-associated mutational processes^9,14^ (**Extended Data** Fig. 9e).

Filtering for the top 50 pseudobulk heteroplasmic variants - selected from a total of 140 variants to focus on those most prominently represented - retained 710 CryoCells (16.5%), which showed an equivalent distribution of states compared to the overall cell type composition (**Fig. 6c,d**). Further, hierarchical clustering of these CryoCells revealed distinct clones, which were predominantly composed of a single cell type. Notably, nearly all SMC-associated clones were defined by a single mtDNA variant, suggesting a predominantly oligoclonal architecture indicative of clonal expansion as intimated in model systems^70,78^ (**Fig. 6e**). Several mtDNA variants exhibited high pseudobulk heteroplasmy with pronounced cell type-specific biases, most prominently in macrophages and fibroblasts (**Fig. 6f,g**). Notably, a subset of fibroblast-like cells harbored clonal mtDNA variants and were distinctly grouped in the UMAP (**Fig. 6h,i** and **Extended Data** Fig. 9**; Methods**). Within this clonally enriched niche, we detected differential chromatin accessibility and transcription factor activity, including elevated *SRF* activity, a transcription factor linked to fibroblast-to- myofibroblast and SMC differentiation^79,80^ (**Fig. 6j**). Together, these findings highlight the utility of mtDNA-based tracing to investigate the clonality in vascular biospecimens and associated cell state transitions otherwise masked by conventional clustering approaches.

## Discussion

Single-cell whole mitochondrial genome sequencing approaches have opened up new avenues for the study of clonality and mitochondrial genetics^8,13,14^. To date, these techniques have revealed a variety of biological insights surrounding the selection of pathogenic mtDNA variants^81–83^ and clonal persistence of innate immune populations^16^. However, their application to solid tissues has remained relatively limited, as each tissue further comes with unique challenges for sampling and tissue dissociation, often resulting in tissue-optimized workflows^17,33^.

Here, we present Cryo-mtscATAC-seq, a robust method for large-scale clonal tracing coupled with cell type annotation which facilitates screening across comprehensively curated data of frozen tissue clinical cohorts. By combining elements of the isolation of cells from the original mtscATAC-seq protocol^8,13–15^ and nuclear preparations strategies from single-nucleus genomics^3^, we established a broadly applicable workflow for isolating CryoCells, enabling multiomic profiling of mitochondrial and nuclear genomes across a wide range of tissues from different organs.

Our protocol is broadly applicable to solid tissues without the need for tissue-specific buffer optimization or specialized equipment. Only minimal input material is required to achieve robust CryoCell recovery, providing a practical alternative to dissociation- based workflows that demand substantial tissue starting material. This might be particularly relevant as clonal expansions in solid tissues often exhibit strong spatial restriction^84^. We demonstrate the versatility of this protocol in *postmortem* human brain tissue, across diverse anatomical regions, revealing local clonal gliogenesis and oligoclonal expansion of microglia. In tumors from opposite ends of the age spectrum - GBM in elderly individuals and neuroblastoma in pediatric patients - we resolve aspects of the clonal architecture, mtDNA copy number heterogeneity and assess changes in cell populations between primary and post-treatment recurrent tumors. Extending to the vascular system, we uncover widespread oligoclonality among smooth muscle cells in the human aorta.

As a CryoCell is not a closed cellular system, we carefully validated which variants can serve as reliable clonal markers, focusing on low-pseudobulk-abundance heteroplasmic variants and additionally applying per-CryoCell VAF thresholds to ensure confident clonal tracing. Typically, putative clonally informative mtDNA variants often exhibited high per-cell VAFs^8,13,14,16^, and recent work has similarly applied binarization with empirically defined thresholds to mitigate ‘mitochondrial’ noise^33^. The implementation of mitoBender further reduces ambient mtDNA contamination, enabling clean datasets comparable to the original mtscATAC-seq protocol^14^. In addition, highly abundant germline homoplasmic variants can be readily exploited for donor demultiplexing to allow for an increase in screening capacity of Cryo-mtscATAC-seq to profile pooled donor samples.

Several limitations should be noted. Mitochondria located within distal cellular branches may be lost during processing (e.g., neuronal axons, dendrites)^45,85^. Consequently, the extrapolation of our protocol to mitochondrial pathologies - where the heteroplasmy of a pathogenic variant within individual mitochondria of a cell may be relevant for pathology has yet to be determined^81,83^. Along these lines, spatial context within the tissue will be critical for understanding cellular interactions, and integrating spatial data with clonal information remains an important goal. Recent studies have demonstrated that mtDNA can be recovered and used as a clonal marker in spatial assays^17,18,86^. However, current approaches are limited by trade-offs between spatial resolution, mitochondrial genomic coverage, and variant identification, or rely on RNA-based readouts, which can be error prone^13^ and may restrict the breadth of clonal information that can be obtained.

In conclusion, Cryo-mtscATAC-seq provides a broadly applicable framework for clonal tracing from frozen clinical specimens, enabling the simultaneous interrogation of mitochondrial and nuclear genomic features across diverse tissues and disease contexts.

## Methods

### Human samples

Human brain tissue samples (ALS and GBM) were obtained from the Brain Bank of Neuropathology, Charité – Universitätsmedizin Berlin. Pathological diagnosis and characterization were performed and confirmed by a certified neuropathologist. GBM samples were obtained from two IDH-wildtype male patients (71 and 61 years). ALS tissue was collected from an 80-year-old ALS–TDP patient (post-mortem interval 12 h) and stored at –80 °C at the Brain Bank of Neuropathology, Charité. Additional ALS tissue from the subventricular zone (SVZ; 63-year-old male, post-mortem interval 20 h) was derived from a donor distinct from that used for the regional analysis. SVZ tissue from a patient with multiple sclerosis (57-year-old male, post-mortem interval 4 h 45 min) was obtained from the Netherlands Brain Bank (NBB). All procedures involving human tissue were conducted in accordance with the Declaration of Helsinki. Autopsy samples were obtained from the Biobank of the Department of Neuropathology, Charité – Universitätsmedizin Berlin. The collection of human brain samples and their subsequent use in this study were approved by the local ethics committee (EA1/368/20, EA1/144/13, EA2/121/20). All donors, or their legal representatives, provided consent for the donation of brain tissue prior to autopsy.

Human neuroblastoma samples were obtained from the Department of Pediatric Oncology and Hematology at Charité - Universitätsmedizin Berlin. Snap-frozen tumor specimens were sampled under the ethical approval EA2/012/23.

Aortic tissue was obtained from a 37-year-old male with a root-dominant thoracic aortic aneurysm (root diameter, 52 mm; proximal ascending, 40 mm; mid-ascending, 38 mm). Sampling was restricted to macroscopically unaffected regions of the proximal and mid-ascending aorta. Tissue procurement and classification were performed by cardiac surgeons at Yale University. Collection of specimens and clinical metadata was approved by the Yale University Institutional Review Board (IRB #2000031944) with a waiver of informed consent. All procedures adhered to institutional ethical guidelines.

### Sample Cryosectioning and Pre-processing

For each sample, 70 - 100 µm cryosections were prepared from fresh-frozen tissue using a cryostat (Feather S35 blades, chamber at –20°C). Sectioning was performed at –18°C, and tissue scrolls were immediately transferred to pre-cooled tubes and stored at –70°C until processing.

### Fixation and CryoCell Isolation

Frozen tissue sections were transferred directly to pre-cooled microcentrifuge tubes and thawed on ice during fixation in formaldehyde (FA) (16% (wt/vol), Thermo Fisher Scientific, cat. no. 28906), diluted in 1× DPBS (ThermoFisherScientific 14190169).

Fixation conditions were tissue-specific: ALS, GBM, NB, and aortic samples were fixed with 1% formaldehyde (FA) for 10 minutes. In some cases, however, a reduced concentration of 0.5% FA was applied for the mixing experiments and selected GBM samples, depending on the experimental batch. The fixed samples were quenched by the addition of 0.125 M glycine (2.5 M; Boston Bioproducts, cat. no. C43755) and gentle inversion and incubation on ice for 5 minutes, followed by centrifugation (400g, 5 min, 4 °C). Pellets were resuspended in chilled isolation buffer (10 mM Tris-HCl pH 7.4, 10 mM NaCl, 3 mM MgCl₂, 1% BSA) and homogenized with a disposable pestle (Fisher Scientific). Suspensions were sequentially passed through 70 μm and 30 μm strainers using a syringe plunger, and nuclei were collected by centrifugation (100g, 10 min, 4 °C). The pellet was resuspended in 1× nuclei buffer and stained with DAPI (1:10,000) for counting. Approximately 8,000 fixed CryoCells were targeted for downstream processing via the modified 10x Genomics-based mtscATAC-seq workflow (see below). Overloading was avoided to reduce debris-associated clogging and doublet rates.

### Library Preparation and Sequencing

CryoCells were processed using the Chromium Next GEM Single Cell ATAC v2 chemistry (10x Genomics) according to the manufacturer’s protocol. Briefly, nuclei were subjected to tagmentation with barcoded transposase and encapsulated into Gel Bead-in-Emulsions (GEMs) using the Chromium Controller X. Linear amplification was performed using a C1000 Touch Thermal Cycler (Bio-Rad), followed by GEM disruption, DNA purification, and indexing PCR (9 cycles). Final libraries were quantified with the Qubit dsDNA High Sensitivity Assay Kit (Invitrogen) and assessed for fragment size distribution on a Bioanalyzer 2100 with High Sensitivity DNA chips (Agilent). Sequencing was carried out at the BIH Genomics Core Facility on Illumina NovaSeq 6000 and NovaSeq X Plus platforms using the following configuration: 100 bp Read 1, 8 bp i7 index, 16 bp i5 index, and 100 bp Read 2.

### Mouse Samples and Cryopreservation

Spleens were harvested from a 25-month-old female *Rosa-CAG-LSL-tdTomato-WPRE* mouse (homozygous for *Rosa*-tdTomato, C57BL/6J background). The spleen was bisected; one half was immediately flash-frozen in 2-methylbutane, and stored at −70 °C. Prior to freezing, tissue was cut into small fragments, rinsed in 1× HBSS, and spatial orientation was documented when relevant. The remaining half was used to generate single-cell suspensions. Tissue was mechanically dissociated through a 40 µm cell strainer (Falcon) in 3–5 ml PBS containing 1% FBS and 2 mM EDTA (PBE). Cell suspensions were filtered (70 µm, Miltenyi), washed with PBE, and centrifuged (400 × g, 5 min, 4 °C). Pellets were resuspended in 3–4 ml ACK lysis buffer, incubated for 4 min at room temperature, quenched with ice-cold PBE, filtered (70 or 100 µm), and centrifuged. Final cell suspensions were resuspended in 5 ml PBE, counted, and cryopreserved in Bambanker medium (Bulldog Bio).

### Mouse splenocytes preparation

Cryopreserved cells were retrieved from liquid nitrogen and thawed rapidly in a 37 °C water bath. Thawed cells were transferred to a 50-ml conical tube, and the cryovial was washed with 1 ml prewarmed RPMI medium supplemented with 10% FBS, which was added dropwise to the tube while gently swirling. Cells were diluted sequentially (1:1) with a prewarmed medium to a total of 50 ml. The suspension was centrifuged (400g, 5 min, 4 °C), the supernatant was discarded, and the pellet was resuspended in 10–25 ml medium depending on cell yield. Cells were washed once more by topping up to 50 ml with medium, centrifuging under the same conditions, and resuspending the pellet in 1 ml FACS buffer (PBS supplemented with 2% FBS and 2 mM EDTA)^21^. Cell number and viability were determined by Trypan Blue exclusion and DAPI staining using light and fluorescence microscopy. For mtscATAC-seq the protocol was followed as described by Lareau et al.^21^. Briefly, cells were pelletted (400g, 5 min, 4 °C), the supernatant was removed, and the pellet was resuspended in 1x DPBS. Suspensions were transferred to 1.5-ml DNA LoBind tubes (Eppendorf) by sequential washing with Fixation was carried out by adding 16% formaldehyde (Thermo Fisher Scientific) to cell suspension in 1x DPBS to a final concentration of 1%, inverting several times, and incubating at room temperature for 10 min. Crosslinking was quenched with 2.5 M glycine (final concentration 0.125 M) by inversion and incubated for 5 min at room temperature. Cells were washed twice with 1 ml ice-cold FACS buffer (1x DPBS supplemented with 2% FBS and 2 mM EDTA), centrifuged (400g, 5 min, 4 °C), and the supernatant was discarded. Library preparation was identical to the CryoCell workflow described above and using the Chromium Next GEM Single Cell ATAC v2 chemistry (10x Genomics) according to the manufacturer’s protocol, as briefly described in Library Preparation and Sequencing.

### Immunofluorescence staining

For cytospin preparations, 200 µl of cell suspension was loaded onto glass slides using a Cytospin 4 cytocentrifuge (Thermo Fisher Scientific; ASHA78300003) at 800 rpm for 8 min (program 8). Cell areas were encircled with a hydrophobic pen, fixed in 4% formaldehyde for 15 min at room temperature, and washed three times in PBS for 5 min each. Blocking was performed for 60 min at room temperature in PBS containing 1% BSA and 10% horse serum. Primary antibodies were diluted in blocking buffer and incubated overnight at 4 °C: rabbit anti-TOMM20 (1:400; Abcam) together with phalloidin-iFluor 568 (1:200; Abcam). Slides were washed three times in PBS and incubated with goat anti-rabbit Alexa Fluor 488 (1:1000; Thermo Fisher Scientific) together with DAPI for 1 h at room temperature. After washing, slides were mounted with PBS, sealed with coverslips using nail polish, and imaged on a Leica DMi8 fluorescence microscope.

### Mouse spleen preprocessing and analysis

Sequencing data were processed with cellranger-atac count (version 2.1.0, 10x Genomics) using alignment to a hard-masked GRCm38 (mm10) reference genome in which nuclear regions homologous to the mitochondrial genome were excluded to minimize misalignment. Mitochondrial single-cell variant calling was carried out with mgatk as described by Lareau et al.^14^ generating variant-level quality metrics including the number of confidently detected cells (n_cells_conf_detected), strand concordance, mean coverage, and the log₁₀-transformed variance-to-mean ratio (VMR). High-confidence variants were retained by applying previously established empirical thresholds for these metrics. Nuclear doublets were identified and removed with Amulet^30^. Each dataset was preprocessed independently in R (version 4.3.1) using the Signac^87^ (version 1.11.0) and Seurat (version 4.4.0) packages. For integration of the different regions, a common peak set was called between the dataset, the datasets were merged and subsetted to high quality cells filtered for each dataset independently. The datasets were integrated using reciprocal LSI anchors and Harmony (version 1.2.3)^88^ batch correction. Dimensionality reduction was performed on corrected embeddings followed by neighborhood graph construction and Leiden clustering. Gene activity scores were computed from ATAC accessibility and log normalized. Annotation was based on defined marker gene expression extracted from the mouse cell atlas^89–91^ and module score calculation.

### MitoBender

For mitoBender, we ran mgatk (mgatk bcall) on the top 20000 barcodes ranked by peak_region_fragments (column 18 in singlecell.csv) and removing barcodes with nonzero mitochondrial count (column 7 in singlecell.csv). We then created input files for cellbender v2 in the legacy 10X mtx format containing the counts for reference and variant alleles in both forward and reverse directions at all variable positions (using either default mgatk filters or a specified list of mtDNA variants). We then ran cellbender v2 on that input, with expected-cells taken from the cellranger output, total-droplets-included by default set at 20000 and low-count-threshold set at 1. We finally replaced values in the raw mgatk output for these variable positions by the cellbender-corrected output. This series of commands is available in mgatk through the mgatk remove-background call (github.com/bihealth/mitobender).

### Preprocessing

Sequencing data were processed with *cellranger-atac count* (version 2.1.0, 10x Genomics) using alignment to a hard-masked GRCh38 reference genome in which nuclear regions homologous to the mitochondrial genome were excluded to minimize misalignment^14^. Mitochondrial single-cell variant calling was carried out with mgatk as described by Lareau et al.^14^, generating variant-level quality metrics including the number of confidently detected cells (n_cells_conf_detected), strand concordance, mean coverage, and the log₁₀-transformed variance-to-mean ratio (VMR). High-confidence variants were retained by applying empirical thresholds on these metrics. Nuclear doublets were identified and removed with Amulet^30^. Each dataset was preprocessed independently in R (version 4.3.1) using the Signac^87^ (version 1.11.0) and Seurat (version 4.4.0) packages. Gene annotations were obtained from Ensembl (EnsDb.Hsapiens.v86) and converted to UCSC-style coordinates. CryocCells were filtered based on chromatin accessibility quality metrics, including transcription start site enrichment (>1.5), nucleosome signal, and nuclear fragment count (>1,000). CryoCells were additionally required to have mitochondrial sequencing depth greater than five reads. CryoCells were further filtered based on their mtDNA depth >5.

### Mixing Experiment

Mixing of nuclear and mtDNA reads between patients at the level of individual CryoCells was assessed using multiple demultiplexing strategies. To capture distinct contamination scenarios, we considered two types of events. First, doublets - either homotypic or heterotypic - were identified using Amulet^30^, representing droplets containing nuclear material from two CryoCells, originating from the same or different patients. Second, to establish the ground truth for CryoCell donor identity, we applied CellSNP-lite^32^ to genotype nuclear variants without a predefined SNP reference to detect common germline SNPs from the 1000 genomes project (using genome1K.phase3.SNP_AF5e2.1toX.hg38.vcf from the cellsnp-lite repo). These genotypes were subsequently used as input for vireo^31^ to infer patient assignments. Given that only two patients were mixed, we expected two distinct nuclear genotypes. Deviations from this - cells assigned as “doublets” by vireo - were interpreted as either true nuclei from two patients or contamination with ambient nuclear material from another CryoCell.

### SVZ specific analysis

For SVZ datasets, we excluded doublets and cells with mtDNA depth ≤5, and integrated vireo^31^ and mtDNA demultiplexing assignments to retain only confidently assigned CryoCells. Chromatin accessibility profiles were processed using TF–IDF normalization followed by dimensionality reduction (SVD, UMAP). mtDNA variants were identified from subsetted mgatk output to define clonal variants per patient. Mean heteroplasmy across filtered variants was computed per donor, aggregated into pseudobulk matrices, and visualized in donor- and variant–ordered heatmaps using ComplexHeatmap. To assess cell type - associated lineage bias, we filtered high-confidence variants, integrated allele frequencies with cell type annotations, applied Kruskal–Wallis tests with Benjamini–Hochberg correction, and summarized mean heteroplasmy and cell counts for bias statistics and visualization.

### ALS specific analysis

Datasets were processed independently as described in preprocessing. For integration of the different regions, a common peak set was called between the datasets. Peak sets from all four regions were converted to GRanges objects and merged into a unified peak set using the reduce function in GenomicRanges. Peaks shorter than 20 bp or longer than 10 kb were excluded, and regions overlapping the ENCODE blacklist were removed. The resulting consensus peak set was used to construct region-specific peak-by-cell count matrices from fragment files. The datasets were merged and subsetted to high quality cells filtered for each dataset independently, furthermore only CryoCells with a mtDNA depth < 100 were retained. The datasets were integrated using reciprocal LSI anchors (dims 1:30) and Harmony^88^ batch correction. Dimensionality reduction was performed on corrected embeddings followed by neighborhood graph construction and Leiden clustering (resolution 0.6). Gene activity scores were computed from ATAC accessibility and log-normalized, and cell-type-specific module scores were derived from curated manually marker gene sets to guide cluster annotation and visualization. Mitochondrial variants were derived from mgatk output^14^, with per-cell allele frequencies calculated from strand-specific counts normalized by coverage. High-confidence variants (n_cells_conf_detected ≥ 5, strand concordance > 0.65, log10[VMR] > −2, mean coverage ≥ 5) were selected from mgatk output per brain region, and region-level mean allele frequencies were calculated. Pseudobulk heteroplasmy profiles were compared across regions. We quantified cell type– associated bias of mitochondrial variants by aligning variant-level allele frequency matrices to grouping cell types and applying Kruskal–Wallis tests with Benjamini - Hochberg correction. To capture clonal representation, CryoCells were considered positive for a variant only if VAF exceeded 20%. Clonal presence per cell type, and summary statistics visualized as −log₁₀-adjusted p-values against group-wise mean heteroplasmy. From the integrated object, we subset CryoCells annotated as Microglia or proinflammatory Microglia. The resulting subset was re-embedded using UMAP computed on Harmony-corrected LSI embeddings (reduction = "harmony", dims = 2:30). A shared nearest-neighbor graph was constructed on the same dimensions, and Leiden clustering was performed with the Louvain variant (algorithm = 3) at resolution 0.4. Gene activity was calculated, and differential peak accessibility was tested on peaks assay (Wilcoxon rank–sum test, min.pct = 0.1, Bonferroni correction). Significant peaks were mapped to closest genes (hg38, Ensembl v86). Cell type biased variants were subsetted to Microglia and proinflammatory Microglia and known artifactual variants (X301A>C, X302A>C, X309C>T, X310T>C, X316G>C, X3109T>C, X189A>G) were removed. For each Region, variants were retained if detected in microglia, with lineage bias ≤ 0.05 and supported by ≥4 cells. Microglia carrying the X1345G>A variant were identified by barcode overlap across regions, visualized in UMAP, and compared to wild-type cells for differential accessibility (Wilcoxon test, min.pct = 0.1), with peaks (|log2FC| > 1.5) linked to nearby genes and visualized at loci such as *HOXA5* using coverage plots

### GBM specific analysis

Datasets were processed independently as described in preprocessing. For integration of the different regions, a common peak set was called between the dataset, the datasets were merged and subsetted to high quality cells filtered for each dataset independently. The datasets were integrated using reciprocal LSI anchors were used to integrate embeddings with IntegrateEmbeddings (dims.to.integrate = 1:30). A UMAP was computed from the integrated embedding with RunUMAP (reduction = "integrated_lsi", dims = 2:30). Graph construction and community detection were performed with FindNeighbors (reduction = "integrated_lsi", dims = 2:30) and FindClusters (Leiden, algorithm = 3, resolution = 0.8). Gene activity scores were calculated from the ATAC accessibility. Annotation of CryoCells was performed using a combination of manual curated canonical marker gene expression and module score calculation based on gene sets derived from Neftel et al.^55^ and Blanco- Carmona et al.^59^. Furthermore large-scale copy number variation profiles were inferred from ATAC-seq signal using EpiAneufinder^92^ to validate overall cluster malignancy. Mitochondrial variants were derived from mgatk output following^14^, with per-cell allele frequencies calculated from strand-specific counts normalized by coverage. High-confidence variants (n_cells_conf_detected ≥ 5, strand concordance > 0.65, log10[VMR] > −2, mean coverage ≥ 5) were selected from mgatk output per tumor. Pseudobulk heteroplasmy profiles were compared primary and recurrence tumor within one patient. To quantify cell type associated bias of mitochondrial variants in the recurrent tumors, we computed variant-level allele frequencies across annotated cell populations for each dataset as described for the human brain data. Differential chromatin accessibility between OPC-like a and malignant OPC-like b populations was assessed using logistic regression on integrated ATAC profiles, followed by motif enrichment analysis (JASPAR2020, version 0.99.10), transcription factor footprinting, and coverage profiling of key regulators (e.g., *SOX2, OLIG2, SOX10, BACH1, JUN, MYT1*) to identify lineage- and malignancy-associated regulatory programs. Furthermore cluster-specific peaks with large effect sizes were mapped to their nearest genes with *ClosestFeature*, and functional annotation was performed by Gene Ontology enrichment analysis. Of note, a subset of CryoCells could not be confidently matched to a defined GBM state and were therefore labeled as undefined tumor cells. These cells were detected across all datasets, displayed elevated EGFR and CD44 gene activity, and, in part, carried characteristic GBM copy-number alterations, including chromosome 7 gain and chromosome 10 loss (**Fig. 4b,d** and **Extended Data** Fig. 5c,e**; Supplementary** Figure 5).

### Neuroblastoma CryoCell Tissue Score

Cryosectioning of each neuroblastoma tumor specimen was performed with a cryostat (CryoStar NX 70) equilibrated to -20°C using S35 Feather microtime blade. Hematoxylin and eosin stain (H&E) staining was performed using standard protocols, followed by microscopic quality assessment to estimate the percentage of intact nuclei (I), necrosis (N), and freezing artifacts (FA). To evaluate tissue integrity and determine suitability for subsequent single-cell experiments, a tissue score (TS) was established based on the microscopic assessment of H&E-stained sections.

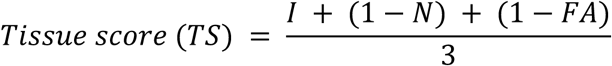

### Neuroblastoma sample analysis

CryoCells were retained if they passed the following thresholds: (nCount_peaks > 250 & nCount_peaks < 20,000, FRiP > 0.1, TSS.enrichment > 1.5, NucleosomeSignal < 2). Cells not meeting these criteria were excluded from downstream analysis. Latent semantic indexing (LSI) was then applied, consisting of term frequency–inverse document frequency (TF–IDF) normalization followed by singular value decomposition (SVD) across all features. Dimensionality reduction and clustering were performed using *RunUMAP* and *FindNeighbors* in Seurat with LSI components 2–30 and resolution = 0.5. Batch correction was carried out with Harmony. Cell type annotation was performed by integrating inferred marker gene activity with label transfer (*FindTransferAnchors*) from single-cell RNA-seq reference datasets^66,93^.

### Aorta specific analysis

In brief, raw Cell Ranger ATAC outputs were loaded into Seurat (version 4.4.0)/Signac (version 1.11.0)^87,94^ and annotated with gene models from EnsDb.Hsapiens.v86. Per-cell quality control metrics were computed, and low-quality cells were excluded if they had fewer than 1,000 detected peaks, TSS enrichment ≤2 or ≥10, or nucleosome signal ≥2. Multiplets were identified using Amulet and removed. Mitochondrial single-cell genotypes were imported from mgatk, and cells with <5× mtDNA depth were excluded. Chromatin accessibility counts were normalized with TF–IDF, dimensionality reduction was performed using SVD and UMAP on LSI components 2–50, and clustering was performed with the Leiden algorithm at a resolution of 0.6. Gene activity scores were computed from chromatin accessibility and log-normalized to support annotation and visualization. Clusters were annotated by manual inspection of gene activity for canonical marker sets curated from Mosquera *et al.*^95^. To isolate a discrete neighborhood in the UMAP, we overclustered the aorta dataset by increasing Leiden resolution to 1.8 (algorithm = 3) on LSI-based embeddings, thereby retaining the targeted UMAP area as a standalone cluster. We then calculated differential accessibility and chromVAR (version 1.24.0) motif activity for the overclustered fibroblast subpopulation (cluster 9; **Extended Data** Fig. 9f) against cluster 13, which under the lower-resolution clustering condition had formed a single aggregate cluster. Mitochondrial DNA variants were imported from mgatk and filtered for high confidence (n_cells_conf_detected ≥ 5, strand correlation > 0.65, log10[VMR] > −2, mean coverage ≥ 5). For the retained variants, pseudobulk allele frequencies were calculated from strand-specific counts normalized by coverage and summarized as mean heteroplasmy per site. To identify clones, only cells carrying ≥20% heteroplasmy in one of the top 50 pseudobulk variants were retained to define per-variant presence/absence. Cells were then clustered using Jaccard (binary) distance with Ward.D linkage. CellBias per variant were identified as described in Quantification of cell type–associated lineage bias of mtDNA variants.

### Use of Artificial Intelligence in manuscript preparation

We enhanced the readability and clarity of this manuscript with assistance from ChatGPT (OpenAI, San Francisco, CA), a large language model artificial intelligence system. ChatGPT was used to check grammar and spelling, improve sentence structure, and overall readability of the text while preserving the original scientific content and conclusions. Suggested revisions were reviewed, edited, and approved by the authors before incorporation into the manuscript. The use of AI assistance was limited to editorial improvements and did not involve generation of research data, statistical analyses, or scientific conclusions. Data collection, analysis, and interpretation were performed by the authors, independent of artificial intelligence.

### Code availability

Detailed descriptions of data analyses and to reproduce the results can be accessed at GitHub (https://github.com/ls-ludwig-lab/Cryo-mtscATAC-seq_reproducibility). The *mitoBender* background correction framework is available as an open-source package at GitHub (github.com/bihealth/mitobender).

### Data availability

Raw sequencing data from human donors and patients have been deposited under controlled access to protect individual genetic privacy at the European Genome-phenome Archive and will be made available upon publication.

## Acknowledgements

We thank the MDC/BIH Genomics Platform, Berlin (FacilityID=1565, The CoreMarketplace: MDC &BIH Technology Platform Genomics) for technical support and sequencing. Computation has been performed on the HPC research cluster of the Berlin Institute of Health. We thank Henrike Scherrer and Stefanie Grosswendt of the Berlin Institute of Health for kindly providing mouse spleen. We are grateful to the members of the Liu and Ludwig labs for valuable discussions. This work was supported by NIH K99/R00 HG012579, R33CA302491 (CAL), UM1 HG012076 (CAL, LSL). LSL is further supported by the Hector Fellow Academy, the Paul Ehrlich Foundation, the EMBO Young Investigator Programme, an Emmy Noether fellowship (LU 2336/2-1) and grants by the German Research Foundation (DFG, LU 2336/3-1, LU 2336/6-1, STA 1586/5-1, LU 2336/8-2, TRR241, SFB1588, Heinz Maier-Leibnitz Award). Individual figures and panels were created with BioRender.com. We are deeply thankful to all patients and their families who agreed to participate in this study.

## Author Contributions

M.S. and L.S.L. conceived and designed the project. M.S. developed the Cryo- mtscATAC-seq protocol, led experimental work, and contributed to data analysis. B.O. developed mitoBender with input from C.A.L. and supervision from D.B.. M.C. provided and analyzed pediatric tumor data. E.F. and F.H. provided ALS samples and discussed data and clinical information. J.C.G., G.C., R.J.B., S.C.S. contributed to protocol development. D.L., P.D., A.H., G.T., R.A., and H.C. provided aorta samples and clinical information; H.C. further discussed data. K.B., M.L., and H.D. provided neuroblastoma samples and clinical information. H.R. provided human brain samples and discussed SVZ data. D.C. and I.L. provided GBM samples and clinical information, with I.L. assisting in the analysis and interpretation of GBM data. S.C.S. initiated the project and contributed to funding acquisition. C.A.L. provided bioinformatics consultation. L.S.L. supervised the project, led protocol development, and provided overall oversight. M.S. and L.S.L. wrote the manuscript with input from all authors.

