## Extended Data Figures for "Cryo-mtscATAC-seq for single-cell mitochondrial DNA genotyping and clonal tracing in archived human tissues"

#### Extended Data Figure 1

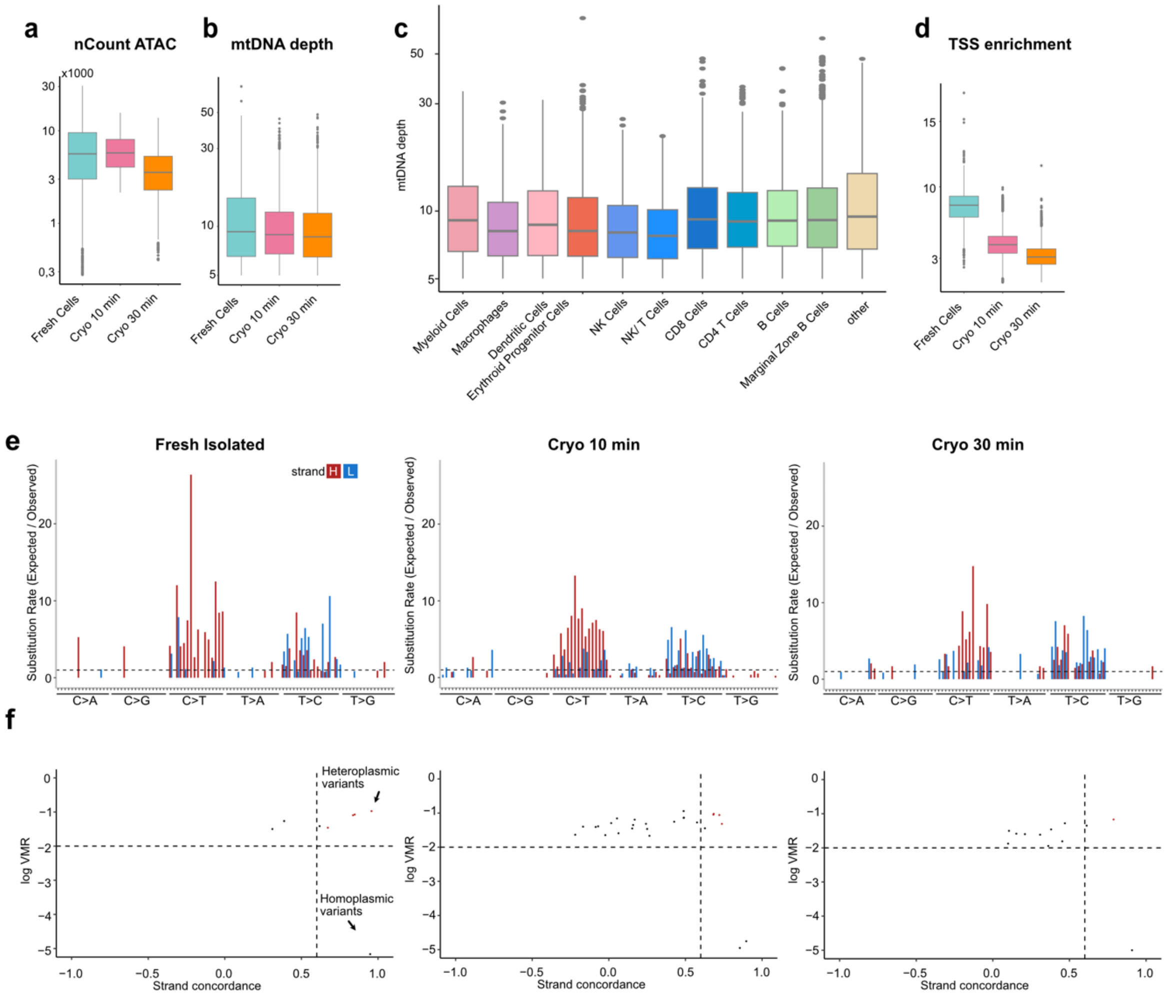


**Extended Data Figure 1 | Quality Measurements of Cryo-mtscATAC-seq in mouse spleen. (a)** Distribution of fragment counts per dataset, illustrating overall sequencing depth across fresh cells and CryoCells fixed for 10 min or 30 min. **(b,c)** mtDNA depth quantified per dataset **(b)** and by celltype **(c),** showing consistent recovery of mtDNA reads across experimental conditions. **(d)** TSS enrichment per dataset. **(e)** Substitution rates (observed/expected) of heteroplasmic mutations identified by mgatk, resolved by mononucleotide and trinucleotide changes and by heavy (H) and light (L) mitochondrial strands. Bar plots represent substitution spectra for fresh cells (left), CryoCells fixed for 10 min (middle), and CryoCells fixed for 30 min (right). **(f)** Identification of high-confidence heteroplasmic variants across conditions. Variants were retained based on strong strand concordance in paired-end sequencing and high variance-to-mean ratio (VMR). Data are shown for fresh cells (left), CryoCells fixed for 10 min (middle), and CryoCells fixed for 30 min (right).

### Extended Data Figure 2

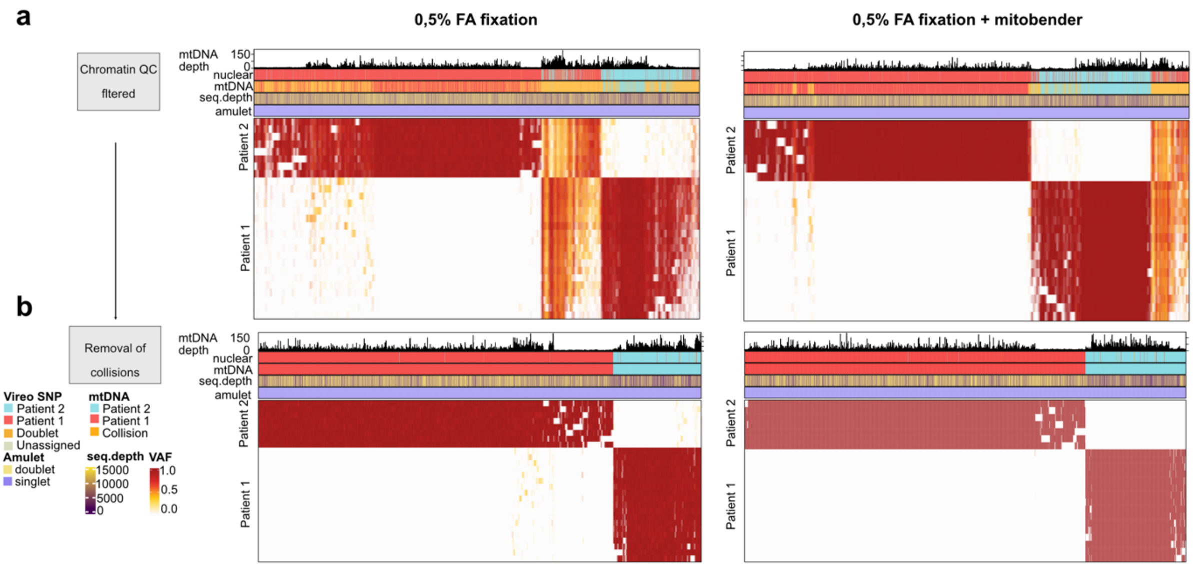


**Extended Data Figure 2 | Heatmap of mtDNA mixing in mixing experiment. (a)** Heatmaps of donor-specific mtDNA variants displaying a bimodal heteroplasmy distribution across cells. Variants were hierarchically clustered, and each column represents a single cell passing chromatin-based QC. Cells are annotated by nuclear genotyping assignment, mtDNA genotyping assignment, Amulet doublet detection, and sequencing depth. Left panel: unfiltered mgatk output. Right panel: mitoBender-filtered matrix. **(b)** Heatmap restricted to cells confidently assigned to patient 1 or patient 2 by their mtDNA profile. Collision cells (with mixed or ambiguous variant signatures) were excluded.

### Extended Data Figure 3

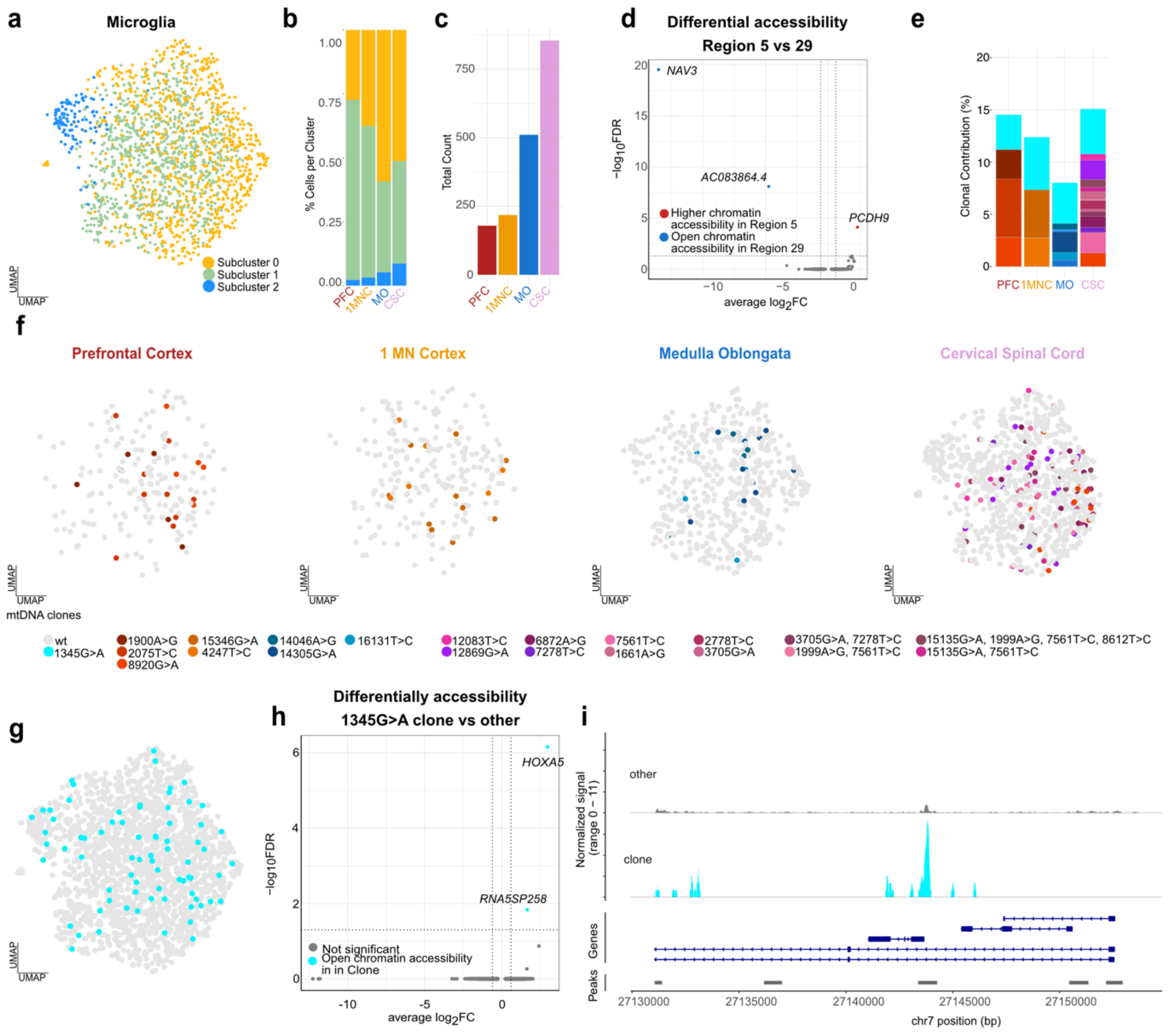


**Extended Data Figure 3 | Oligoclonal microglia in different brain region of an ALS patient. (a)** Subclustering of microglia indicative of distinct chromatin states. **(b)** Proportion of microglia subclusters across anatomical regions. **(c)** Total number of microglia recovered per region. **(d)** Differential chromatin accessibility between microglia from PFC and CSC, shown as average $\log_2$ fold change (x-axis) versus –$\log_{10}$ FDR (y-axis). Genes with region-specific accessibility include *PCDH9* (PFC) and *NAV3* (CSC). **(e)** Clonal composition of microglia across regions, based on mtDNA variants defining individual clones. **(f)** UMAP representation of microglial clones per region (left to right: PFC → CSC). Clones are colored by mtDNA variant identity; CryoCells are assigned if carrying ≥20% VAF of the respective variant. **(g)** UMAP projection of the mt.1345G>A clone spanning all regions. **(h)** Differential chromatin accessibility between the mt.1345G>A clone and all other microglia, revealing higher accessibility in the clone. **(i)** Chromatin accessibility tracks at the *HOXA5* locus, comparing the mt.1345G>A clone with all other microglia.

### Extended Data Figure 4

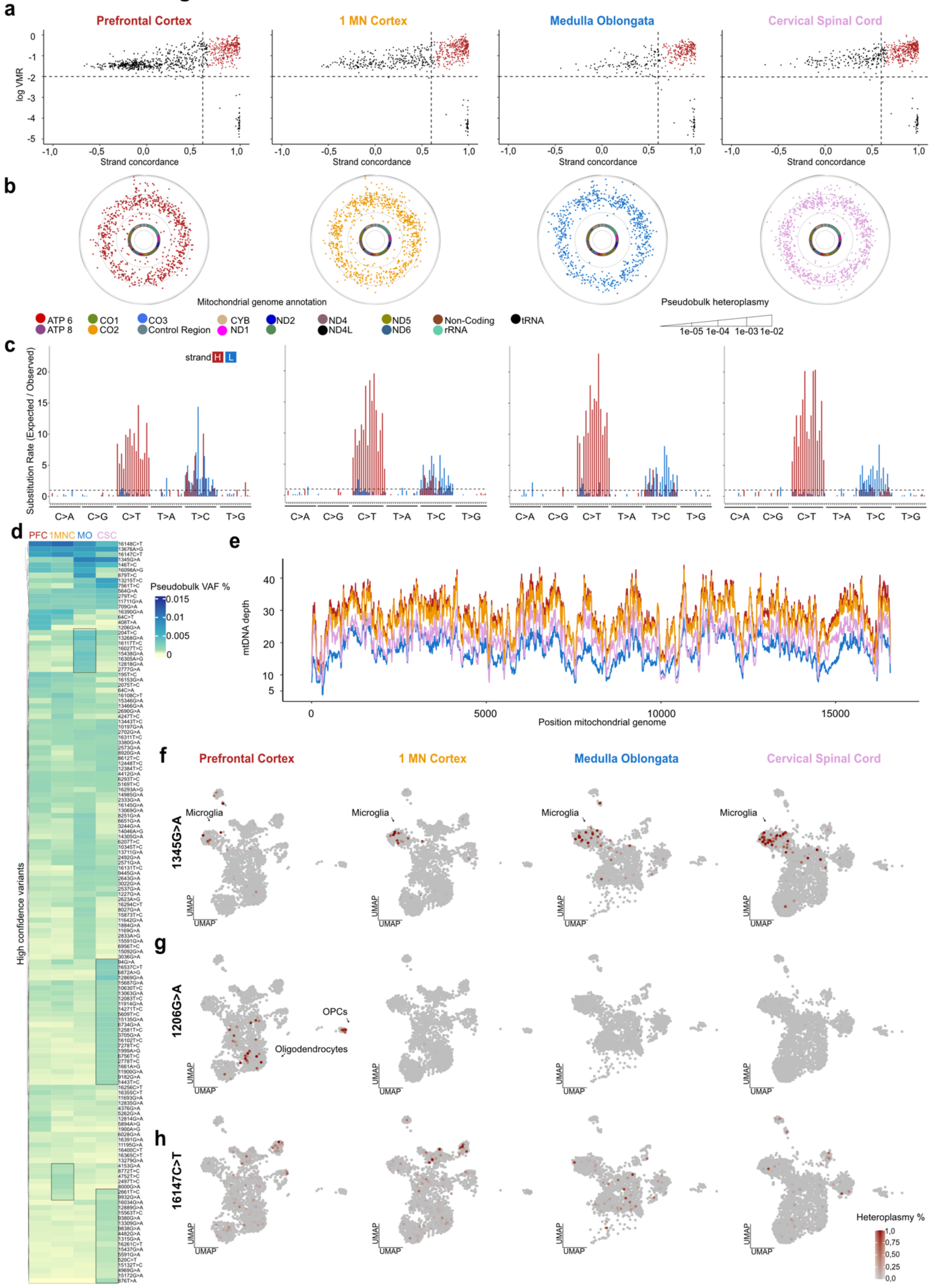


**Extended Data Figure 4 | Mitochondrial genotyping across different brain regions of an ALS patient. (a)** Identification of high-confidence heteroplasmic variants across regions. Variants were retained based on high strand concordance in paired-end sequencing and elevated variance-to-mean ratio (VMR). Shown from left to right: PFC → CSC. **(b)** Circular plots of mgatk-nominated mutations along the mitochondrial genome for each region. The inner circle denotes the mitochondrial genome, color-coded by gene annotation. Each dot represents a variant, positioned radially by % heteroplasmy (inner to outer), with the outer ring representing the heteroplasmy axis. **(c)** Substitution spectra of heteroplasmic mutations (observed/expected), resolved by mononucleotide and trinucleotide changes and by heavy (H) and light (L) strands of the mitochondrial genome, shown for each region (left to right: PFC → CSC). **(d)** Heatmap of high-confidence heteroplasmic variants per region (filters: n_cells_conf_detected ≥ 5, strand_correlation > 0.65, $\log_{10}$(VMR) > –2, mean_coverage ≥ 5). **(e)** Per-nucleotide sequencing depth across the mitochondrial genome for each dataset. **(f–h)** UMAP projections showing per-cell heteroplasmy (%) for representative mtDNA variants across regions (left to right: PFC → CSC). Variants include (f) mt.1345G>A, (g) mt.1206G>A, and (h) mt.16147T>C. Cells are colored by allele frequency, and clones were assigned based on >20% VAF for downstream analyses (see Fig. 2).

### Extended Data Figure 5

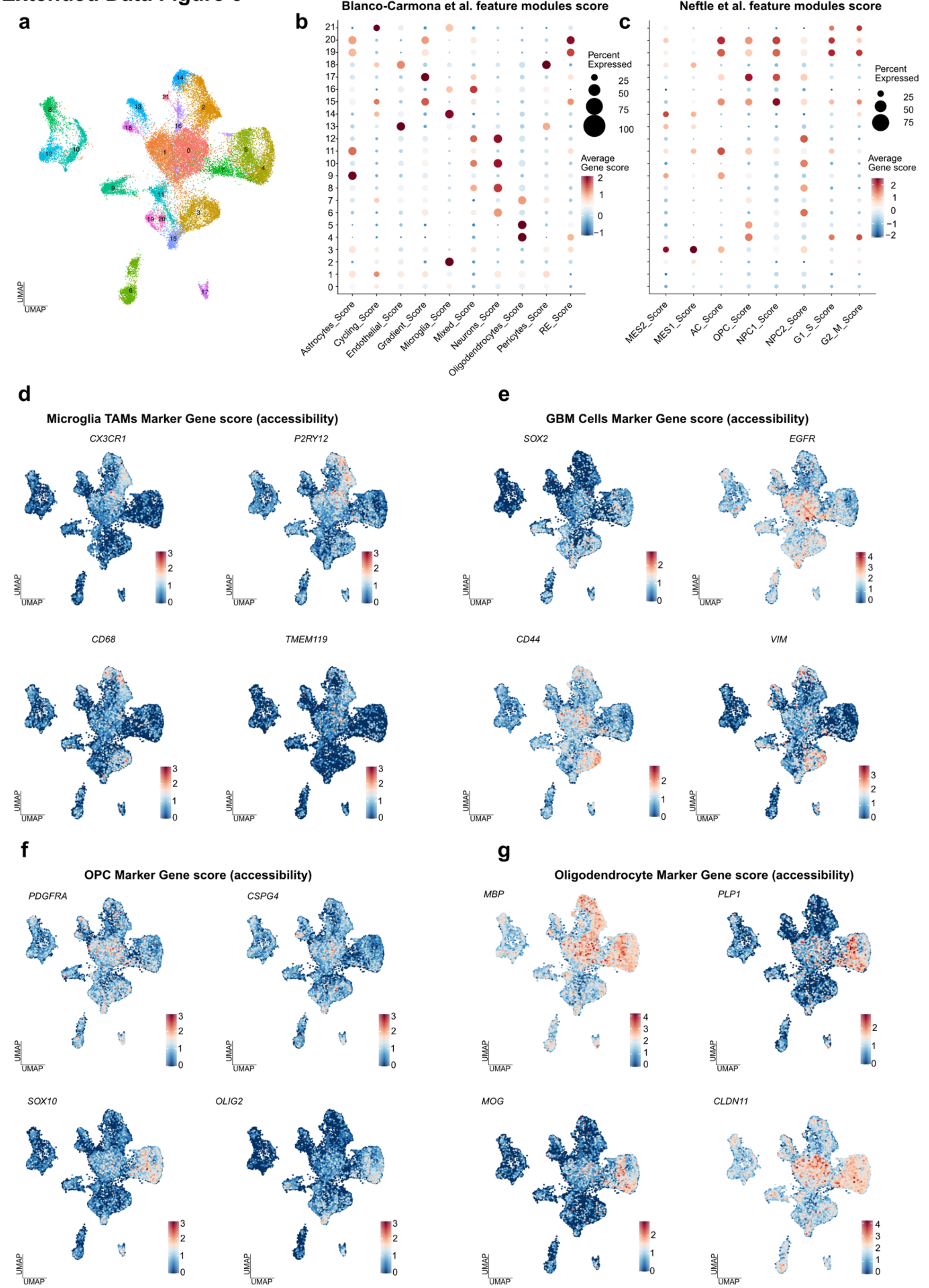


**Extended Data Figure 5 | Clustering and annotation of longitudinal GBM samples. (a)** UMAP embedding of integrated GBM CryoCells clustered by chromatin accessibility. **(b,c)** Dot plots of gene module activity derived from Blanco-Cámara *et*

*al.* and Neftel *et al.*. The x-axis indicates module assignments for each cell type, and the y-axis represents clusters within the integrated dataset. Each dot corresponds to pseudobulk gene activity inferred from chromatin accessibility, with color intensity reflecting average expression and dot size indicating the percentage of cells with detectable signal. **(d–g)** UMAP projections of inferred expression for selected marker genes, highlighting major cellular populations. **(d)** Microglia and tumor-associated macrophage (TAM) markers (*CX3CR1, P2RY12, CD68, TMEM119*). **(e)** GBM-associated markers (*SOX2, EGFR, CD44, VIM*). **(f)** Oligodendrocyte progenitor cell (OPC) markers (*PDGFRA, CSPG4, SOX10, OLIG2*). **(g)** Mature oligodendrocyte markers (*MBP, PLP1, MOG, CLDN11*).

### Extended Data Figure 6

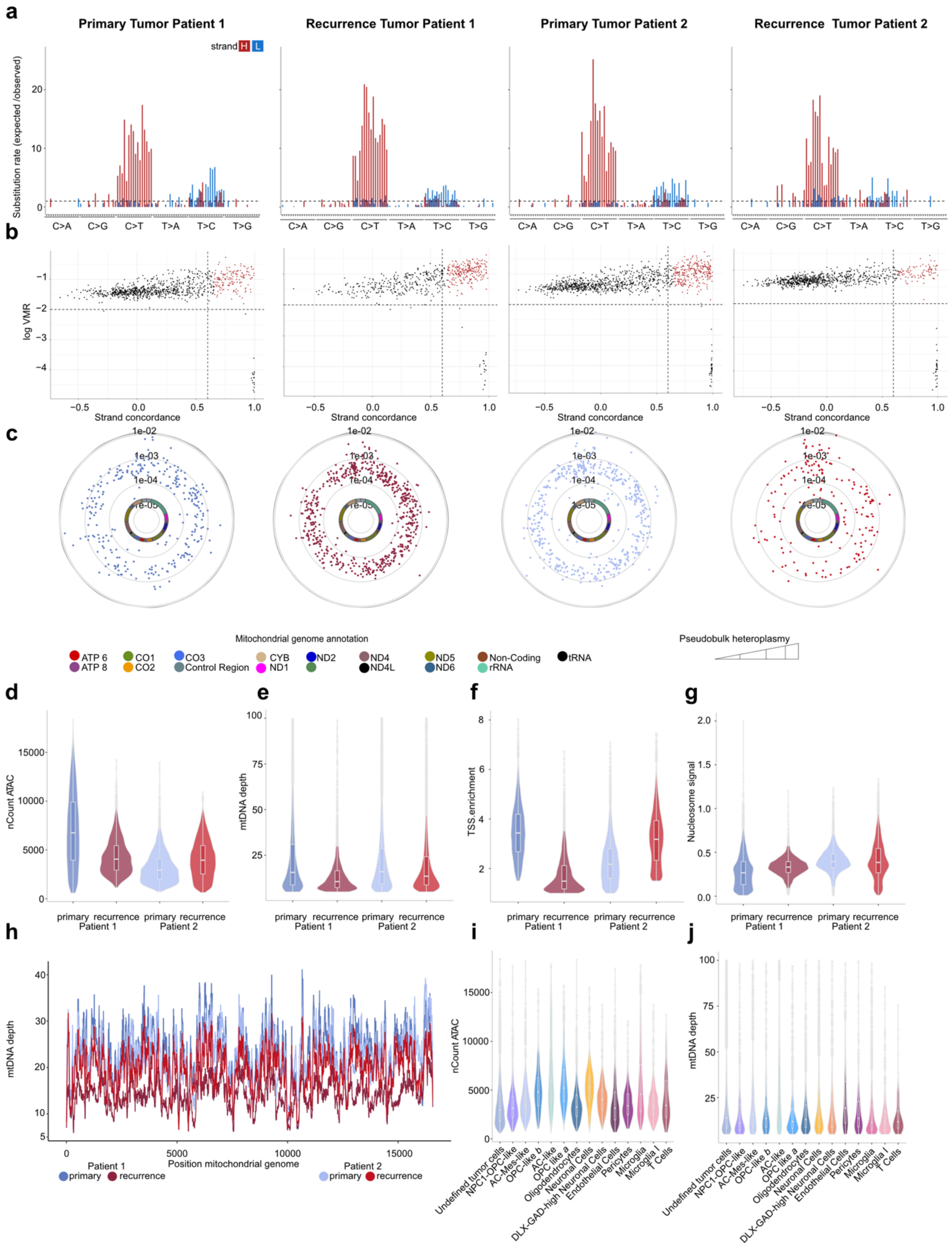


**Extended Data Figure 6 | Mitochondrial genotyping in longitudinal GBM samples. (a)** Substitution spectra of heteroplasmic mutations (observed/expected), resolved by mononucleotide and trinucleotide changes and by heavy (H) and light (L)

strands of the mitochondrial genome. Shown for each dataset from left to right: Patient 1 primary → Patient 2 recurrence. **(b)** Identification of high-confidence heteroplasmic variants across datasets. Variants were retained based on high strand concordance in paired-end sequencing and elevated variance-to-mean ratio (VMR). Shown from left to right: Patient 1 primary → Patient 2 recurrence.**(c)** Circular plots of mgatk-nominated mutations along the mitochondrial genome for each dataset. The inner circle denotes the mitochondrial genome, color-coded by gene annotation. Each dot represents a variant, positioned radially by % heteroplasmy (inner to outer), with the outer ring representing the heteroplasmy axis. Shown from left to right: Patient 1 primary → Patient 2 recurrence. **(d–g)** Standard QC metrics across datasets: **(d)** sequencing depth (nCount ATAC), **(e)** mitochondrial read depth, **(f)** TSS enrichment, and **(g)** nucleosomal signal. **(h)** Per-nucleotide sequencing depth across the mitochondrial genome for each dataset. **(i,j)** Sequencing depth (nCount ATAC) **(i)** and mitochondrial read depth **(j)** stratified by cell type.

### Extended Data Figure 7

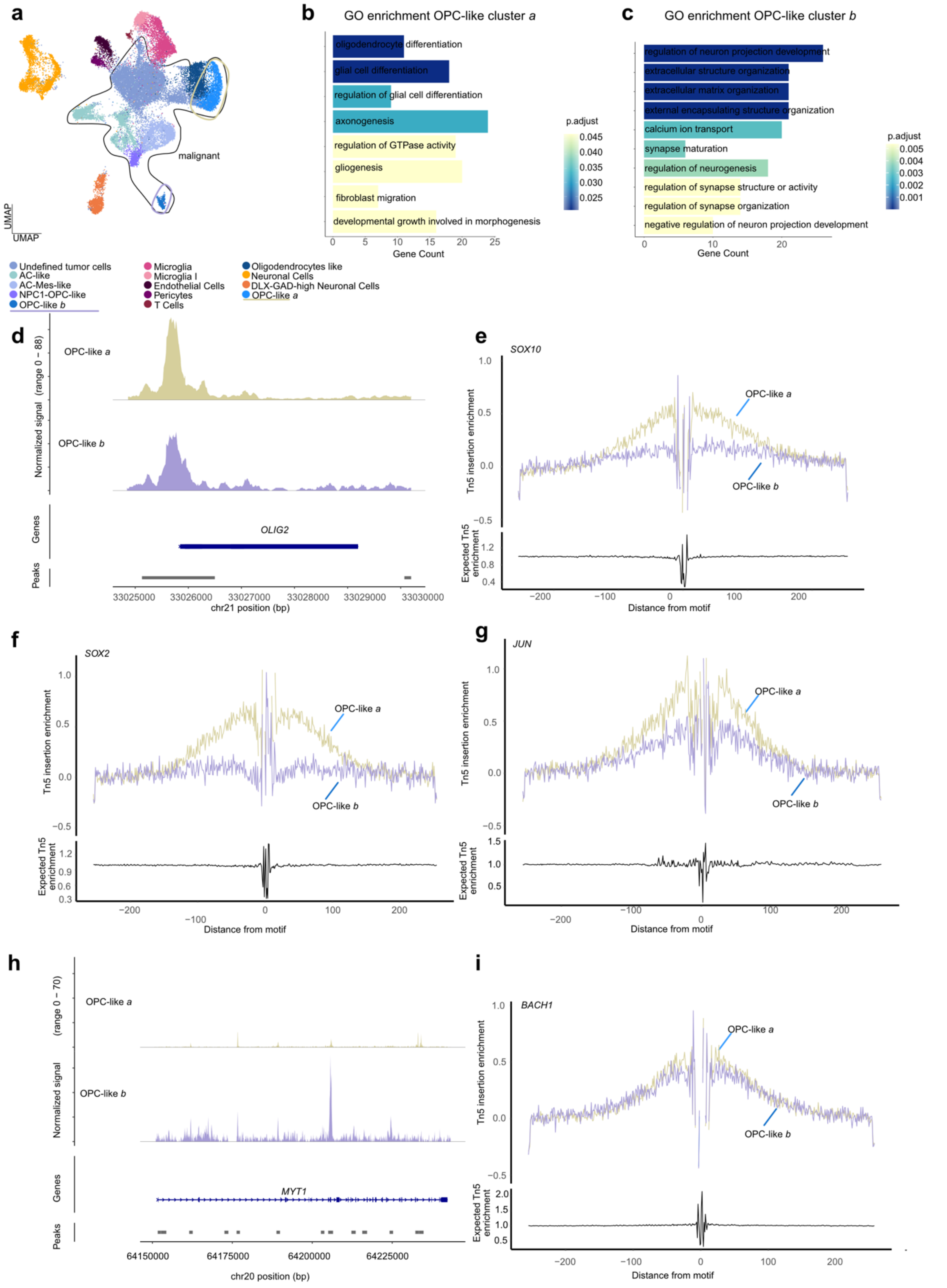


**Extended Data Figure 7 | Characterization of chromatin accessibility in OPC-like cell clusters in GBM. (a)** UMAP embedding of integrated GBM CryoCells colored by cell type. Malignant cells are highlighted, with the OPC-like cluster further separated to distinguish subpopulations. **(b,c)** Gene ontology (GO) enrichment analysis of cellular programs in OPC-like (b) and aggressive OPC-like (c) CryoCells. **(d–i)** Differential chromatin accessibility and transcription factor activity analyses distinguishing OPC-like versus aggressive OPC-like states. **(d)** Chromatin accessibility track at *OLIG2*. **(e)** Transcription factor footprinting for SOX10. (f) Transcription factor footprinting for SOX2. **(g)** Transcription factor footprinting for JUN. **(h)** Chromatin accessibility track at *MYT1*. **(i)** Transcription factor footprinting for BACH1.

### Extended Data Figure 8

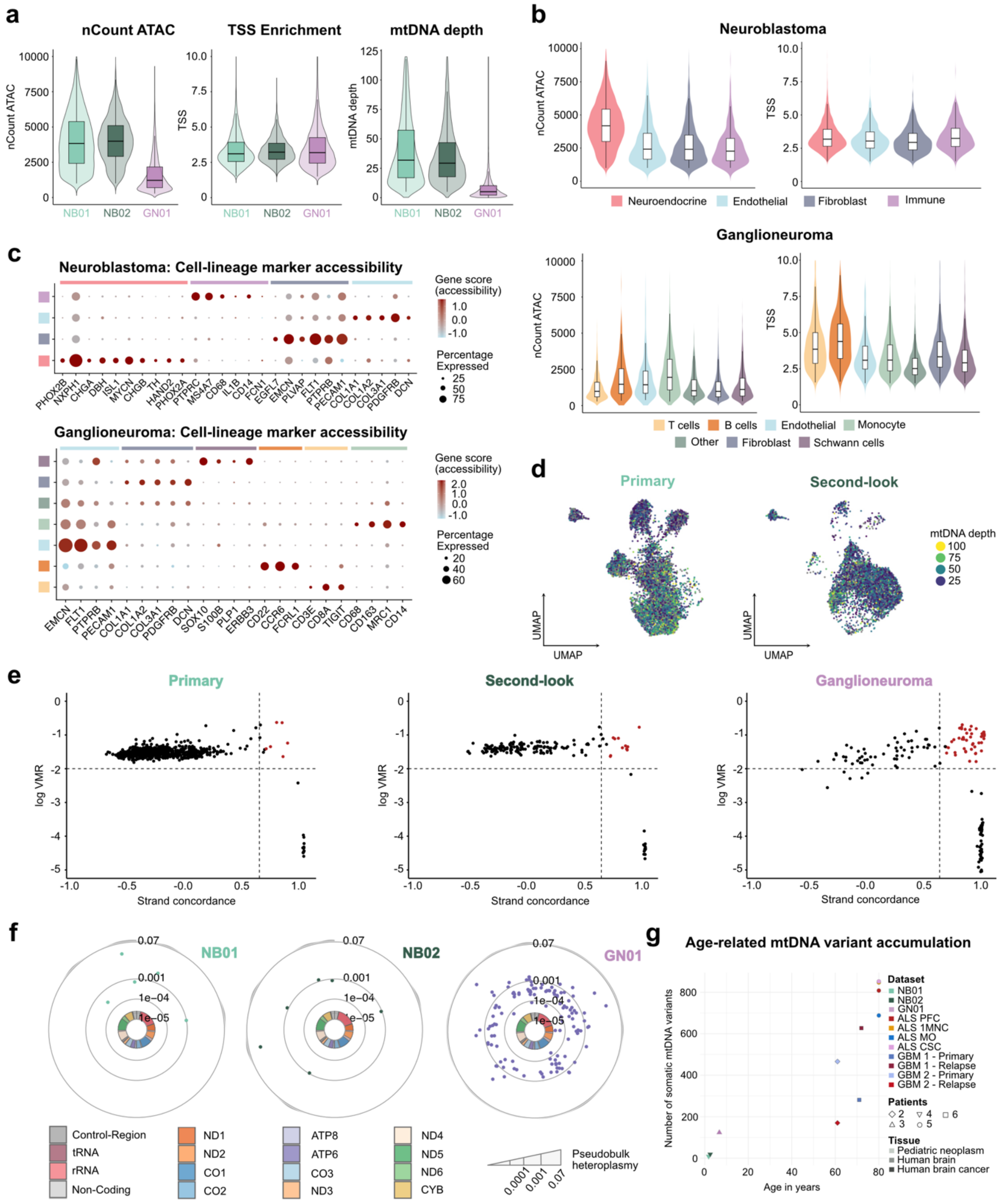


**Extended Data Figure 8 | Cryo-mtscATAC-seq quality metrics and annotation in pediatric neoplasms and age related mtDNA mutation accumulation. (a)** Quality measurements for pediatric neoplasm datasets: sequencing depth (nCount ATAC, left), TSS enrichment (middle), and mtDNA depth (right). NB01 = primary neuroblastoma, NB02 = second-look neuroblastoma from the same patient, GN01 = ganglioneuroma. **(b)** nCount ATAC and TSS enrichment stratified by cell type annotation in integrated neuroblastoma (top) and ganglioneuroma (bottom). **(c)** Gene activity scores of canonical marker genes used for cell type annotation in neuroblastoma (top) and ganglioneuroma (bottom). **(d)** mtDNA depth projected onto UMAP embeddings, shown for neuroblastoma (left) and ganglioneuroma (right). **(e)** Identification of high-confidence heteroplasmic variants across datasets, retained based on strong strand concordance in paired-end sequencing and elevated variance-to-mean ratio (VMR). Shown left to right: NB01 primary neuroblastoma, NB02 second-look neuroblastoma, GN01 ganglioneuroma. **(f)** Circular plots of mgatk-nominated mutations across the mitochondrial genome for each dataset. The inner circle represents the mitochondrial genome color-coded by gene annotation. Each dot corresponds to a variant, positioned radially by % heteroplasmy (inner to outer), with the outer ring representing the heteroplasmy axis. Shown left to right: NB01, NB02, GN01. **(g)** Age-related accumulation of mtDNA variants in human clinical specimens spanning pediatric neoplasms, GBM, and ALS.

### Extended Data Figure 9

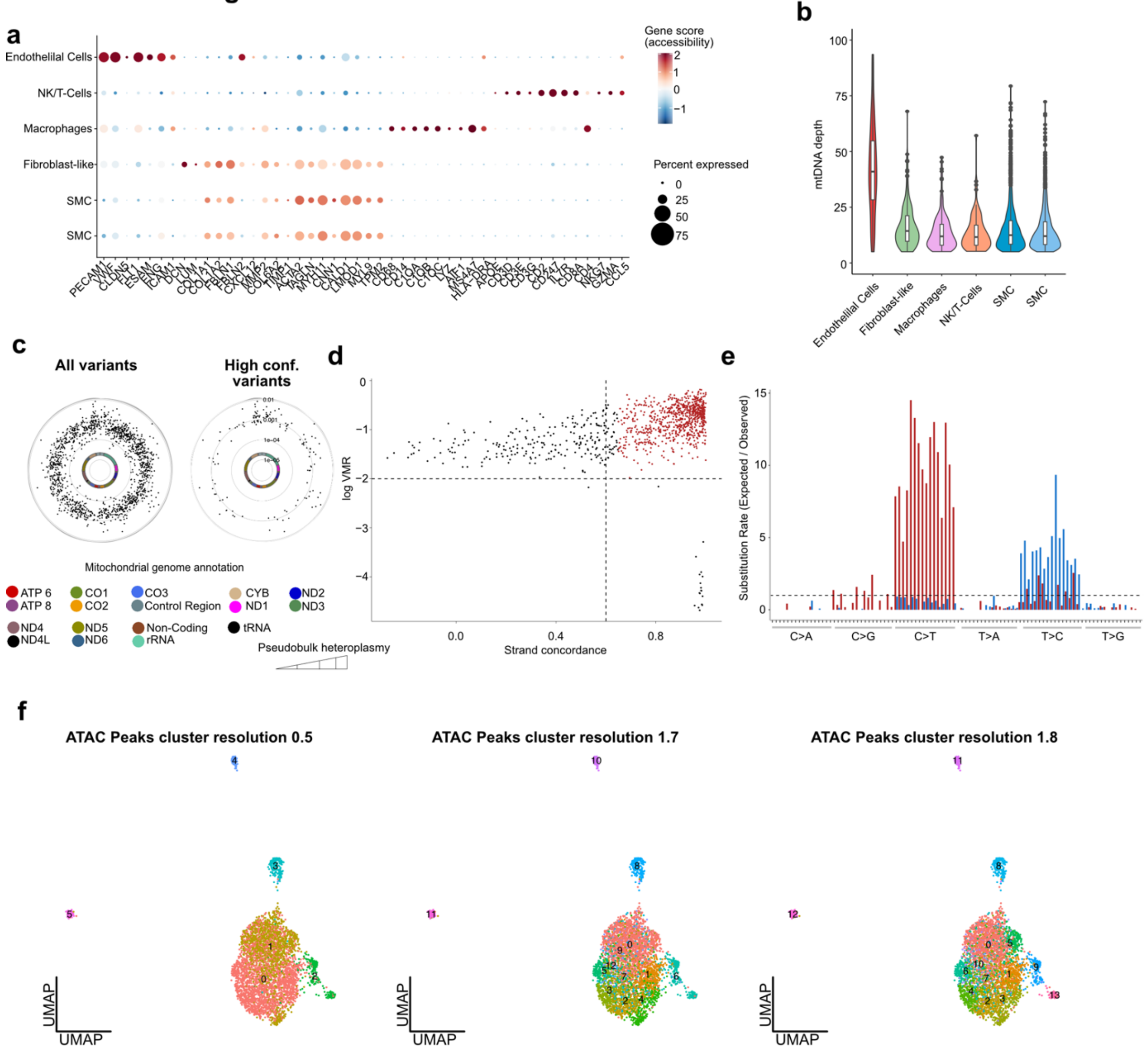


**Extended Data Figure 9 | Human aorta annotation and mitochondrial genotyping. (a)** Dot plot of manually curated marker genes for vascular cell populations within the human aorta. **(b)** mtDNA depth stratified by cell type. **(c)** Circular plots of mgatk-nominated mutations across the mitochondrial genome. The inner circle represents the mitochondrial genome, color-coded by gene annotation. Each dot corresponds to a variant, positioned radially by % heteroplasmy (inner to outer), with the outer ring representing the heteroplasmy axis. Left: all variants identified by mgatk. Right: subset of high-confidence variants (n_cells_conf_detected ≥ 5, strand_correlation > 0.65, $\log_{10}$(VMR) > –2, mean_coverage ≥ 5). **(d)** Identification of high-confidence heteroplasmic variants across datasets, retained based on strong strand concordance in paired-end sequencing and elevated variance-to-mean ratio (VMR). **(e)** Substitution spectra of heteroplasmic mutations (observed/expected), resolved by mononucleotide and trinucleotide changes and by heavy (H) and light (L)

mitochondrial strands. **(f)** UMAP embeddings of clustering based on ATAC chromatin peaks at different resolutions (left: 0.5; middle: 1.7; right: 1.8). At resolution 1.8, cluster 9 corresponds to a clonally enriched niche of fibroblast-like cells.
