## Supplementary Figures for "Cryo-mtscATAC-seq for single-cell mitochondrial DNA genotyping and clonal tracing in archived human tissues"

### Supplementary Figure

#### Supplementary Figure 1

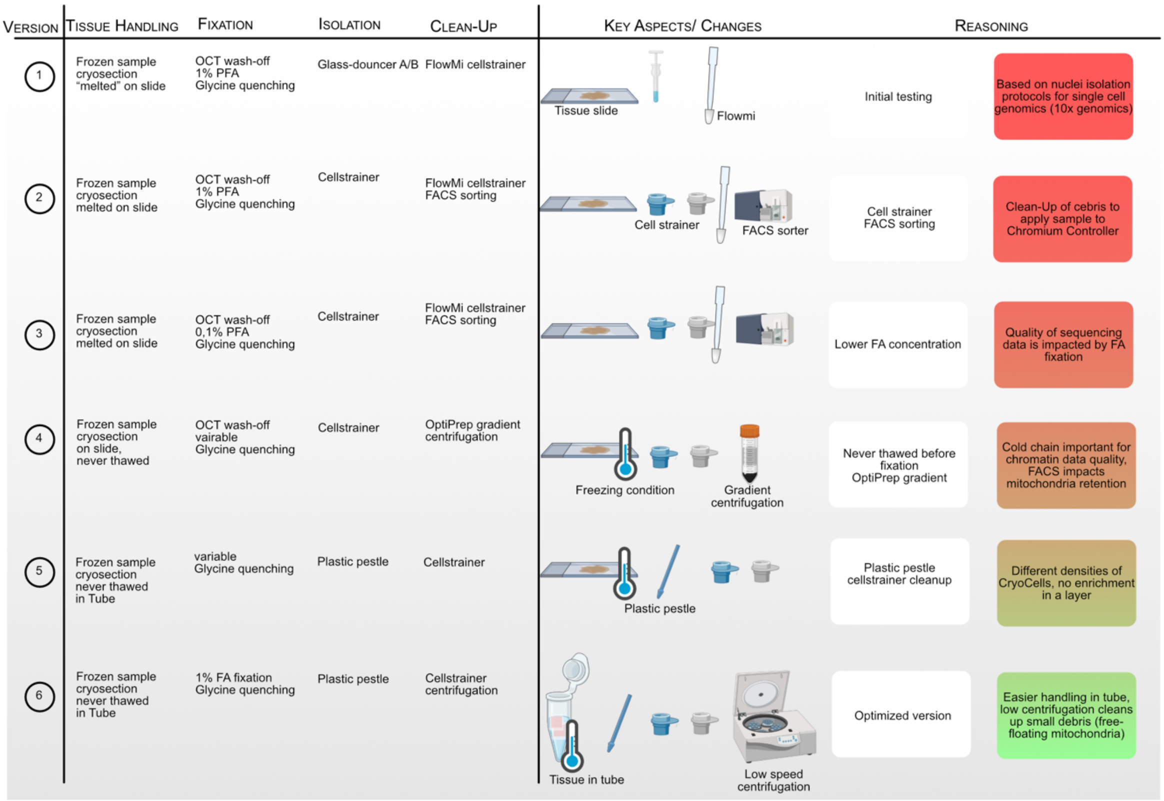


**Supplementary Figure 1| Wetlab protocol evolution for Cryo-mtscATAC-seq.** Overview of establishing the wet-lab protocol, where different isolation methods were systematically tested as indicated until the final workflow was defined.

#### Supplementary Figure 2

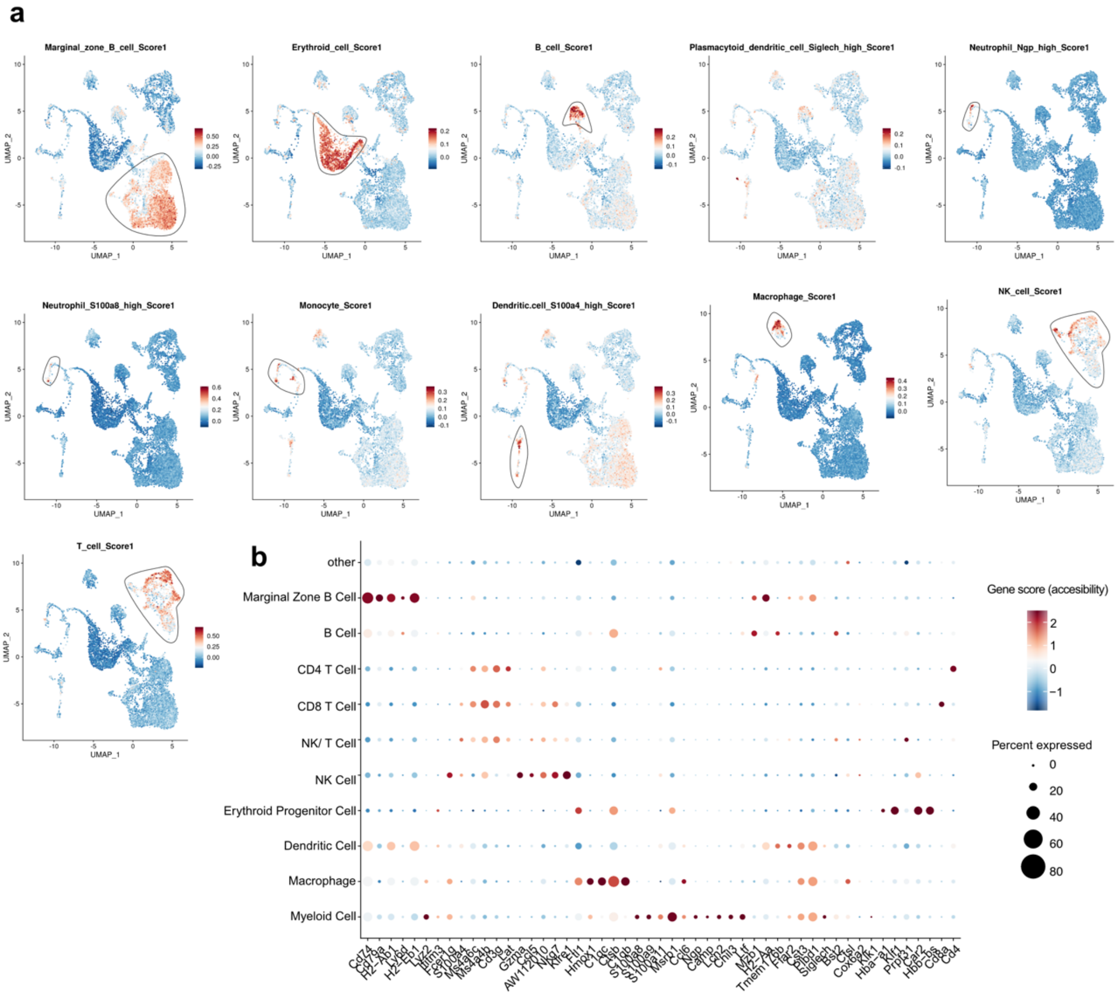


**Supplementary Figure 2| Annotation of mouse spleen. (a)** UMAP representation of manually curated module scores for spleen-resident cell types. Upper panel (left to right): marginal zone B cells, erythroid progenitor cells, B cells, plasmacytoid dendritic cells, neutrophils. Middle panel (left to right): neutrophil (*S100ab*), monocytes, dendritic cells, macrophages, NK cells. Lower panel: T cells. **(b)** Dot plot of manually curated marker genes used for annotation of hematopoietic cell populations.

#### Supplementary Figure 3

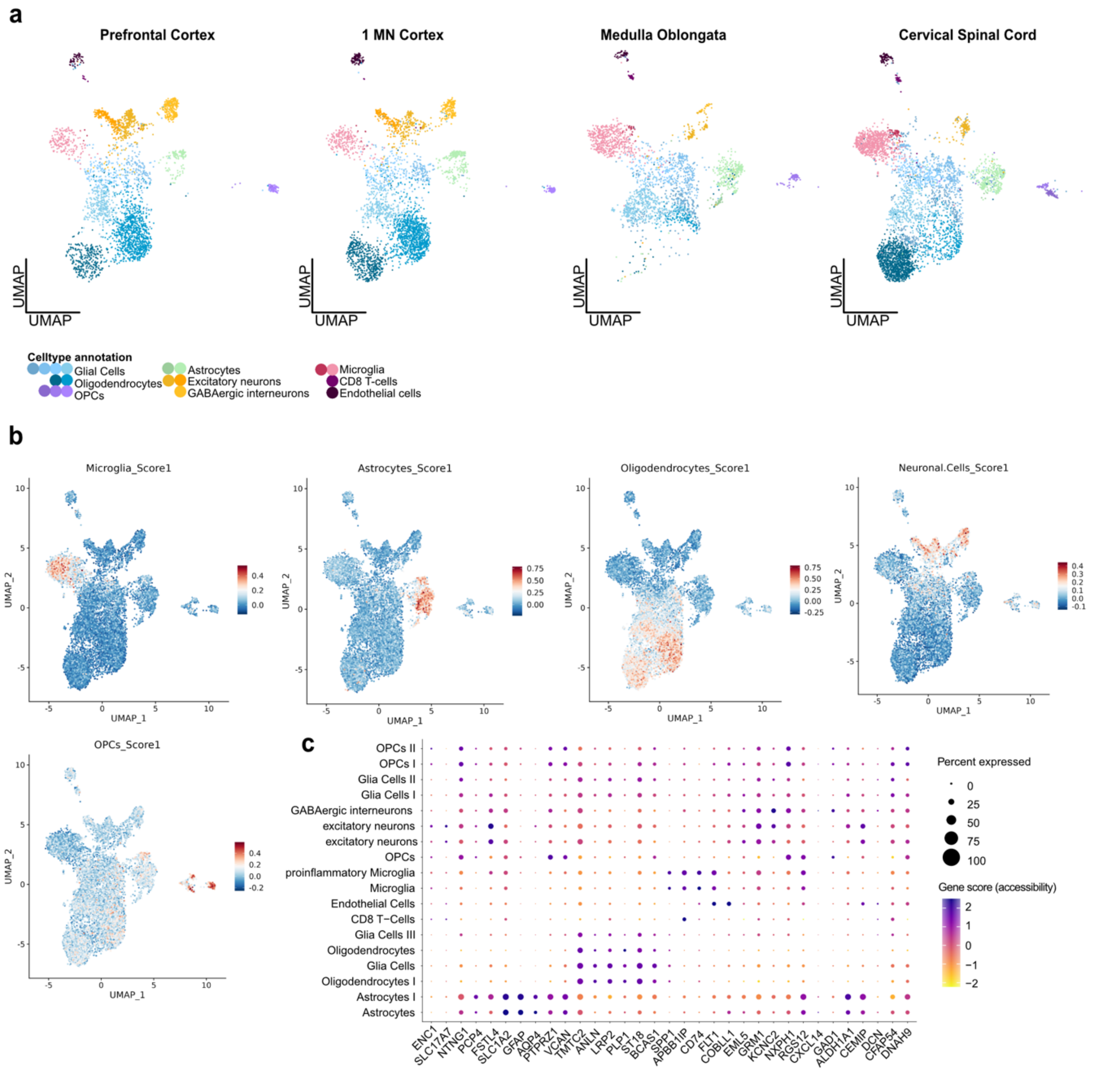


**Supplementary Figure 3 | Annotation of human ALS brain. (a)** UMAP representation of CryoCells divided by anatomical region. **(b)** UMAP representation of manually curated cell type scores. Upper panel (left to right): microglia, astrocytes, oligodendrocytes, neuronal cells, oligodendrocyte progenitor cells (OPCs). **(c)** Dot plot of marker genes used for annotation of major CNS cell types.

#### Supplementary Figure 4

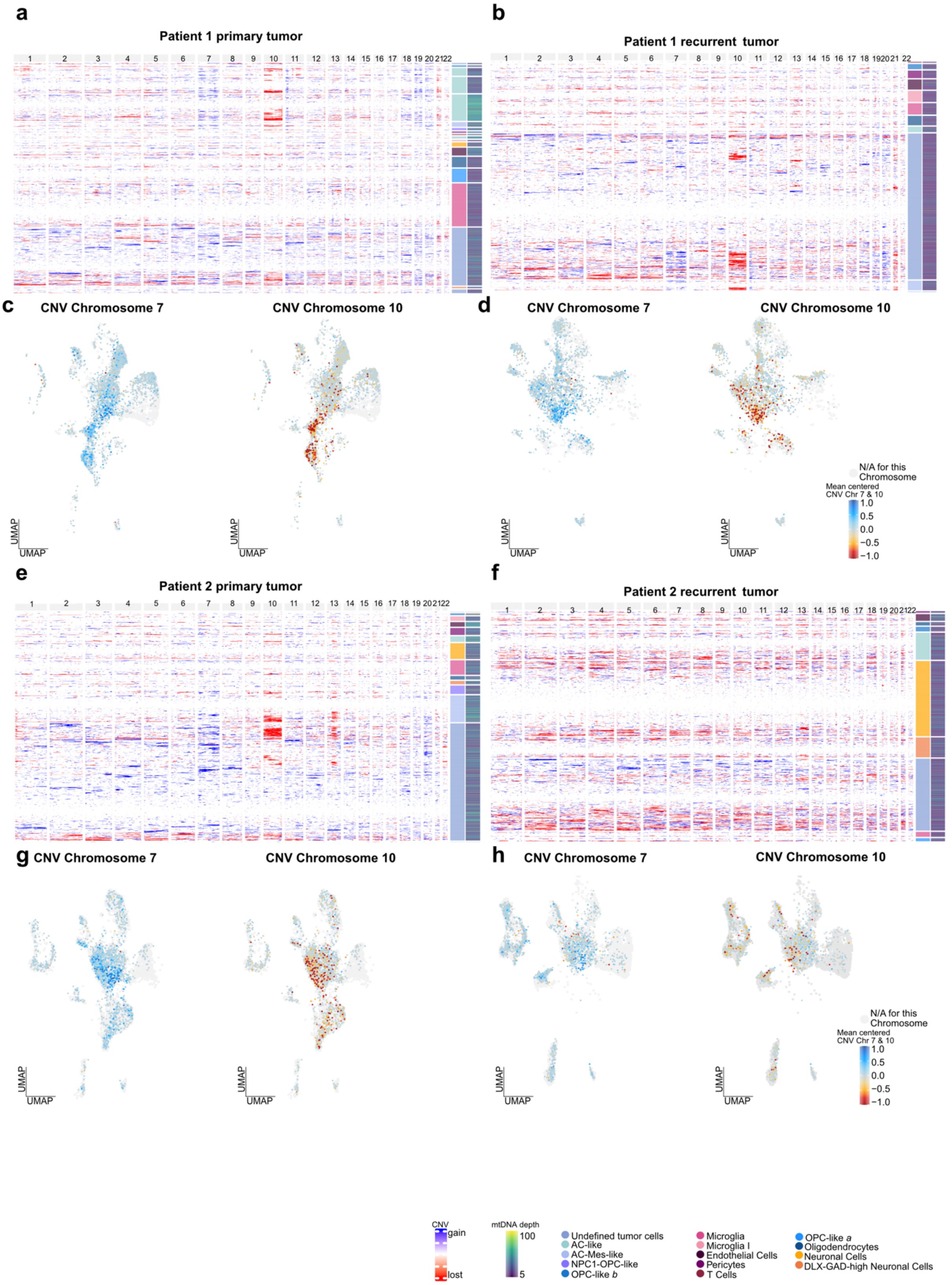


**Supplementary Figure 4| Copy number variant analysis of GBM samples. (a,b)** Karyograms of copy-number variations (CNVs) for patient 1 (**a**, primary; **b**, recurrence). **(c,d)** UMAP projections of mean-centered CNVs for chromosome 7 (left)

and chromosome 10 (right) in patient 1 **(c,** primary; **d,** recurrence). **(e,f)** Karyograms of CNVs for patient 2 (**e**, primary; **f**, recurrence). **(g,h)** UMAP projections of mean-centered CNVs for chromosome 7 (left) and chromosome 10 (right) in patient 2 (**g**, primary; **h**, recurrence).

#### Supplementary Figure 5

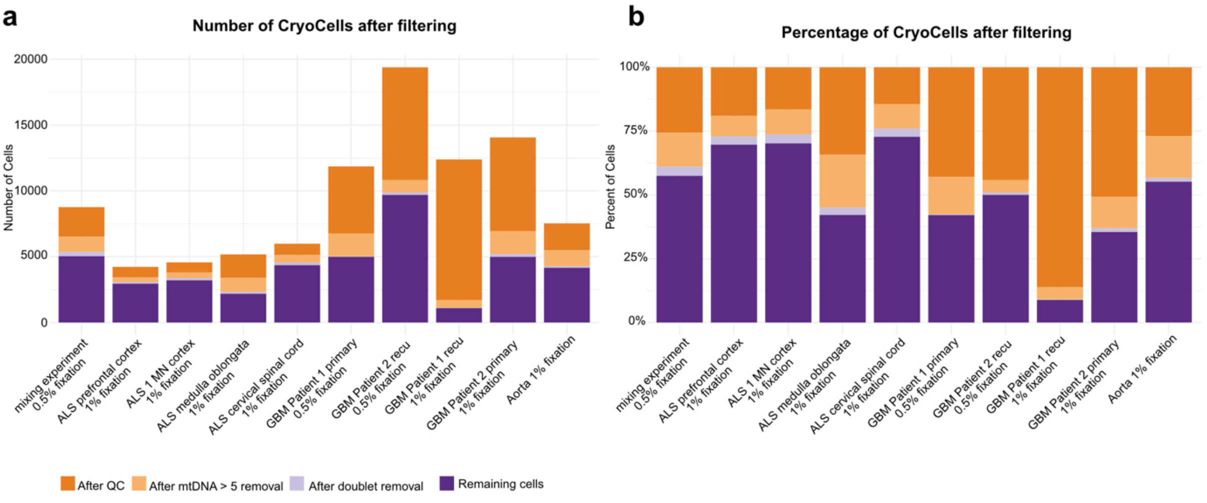


**Supplementary Figure 5 | CryoCell dropout after filtering. (a)** Number of CryoCells retained at successive filtering steps for each human dataset analyzed in this study. Bars indicate cells removed after chromatin accessibility QC filtering (orange), additional filtering for mitochondrial DNA depth >5 (bright orange), and removal of nuclear doublets detected with Amulet (light purple). Remaining cells carried forward for downstream analysis are shown in purple. (**b)** Proportional representation of CryoCells at each filtering step, displayed as percentages relative to the initial number of captured CryoCells.
